# Inhibition of RhoA-mediated secretory autophagy in megakaryocytes mitigates myelofibrosis in mice

**DOI:** 10.1101/2024.12.04.626665

**Authors:** Isabelle C. Becker, Maria N. Barrachina, Joshua Lykins, Virginia Camacho, Andrew P. Stone, Bernadette A. Chua, Robert A.J. Signer, Kellie R. Machlus, Sidney W. Whiteheart, Harvey G. Roweth, Joseph E. Italiano

## Abstract

Megakaryocytes (MKs) are large, polyploid cells that contribute to bone marrow homeostasis through the secretion of cytokines such as transforming growth factor β1 (TGFβ1). During neoplastic transformation, immature MKs accumulate in the bone marrow where they induce fibrotic remodeling ultimately resulting in myelofibrosis. Current treatment strategies aim to prevent MK hyperproliferation, however, little is understood about the potential of targeting dysregulated cytokine secretion from neoplastic MKs as a novel therapeutic avenue. Unconventional secretion of TGFβ1 as well as interleukin 1β (IL1β) via secretory autophagy occurs in cells other than MKs, which prompted us to investigate whether similar mechanisms are utilized by MKs. Here, we identified that TGFβ1 strongly co-localized with the autophagy marker light chain 3B in native MKs. Disrupting secretory autophagy by inhibiting the small GTPase RhoA or its downstream effector Rho kinase (ROCK) markedly reduced TGFβ1 and IL1β secretion *in vitro*. *In vivo*, conditional deletion of the essential autophagy gene *Atg5* from the hematopoietic system limited megakaryocytosis and aberrant cytokine secretion in an MPL^W515L^-driven transplant model. Similarly, mice with a selective deletion of *Rhoa* from the MK and platelet lineage were protected from progressive fibrosis. Finally, disease hallmarks in MPL^W515L^-transplanted mice were attenuated upon treatment with the autophagy inhibitor hydroxychloroquine or the ROCK inhibitor Y27632, either as monotherapy or in combination with the JAK2 inhibitor ruxolitinib. Overall, our data indicate that aberrant cytokine secretion is dependent on secretory autophagy downstream of RhoA, targeting of which represents a novel therapeutic avenue in the treatment of myelofibrosis.

**One Sentence Summary:** TGFβ1 is released from megakaryocytes via RhoA-mediated secretory autophagy, and targeting this process can alleviate fibrosis progression in a preclinical mouse model of myelofibrosis.

## Introduction

Megakaryocytes (MKs) are large, polyploid cells that primarily reside within the bone marrow (*1*). They differentiate from hematopoietic stem cells (HSCs) upon binding of the growth factor thrombopoietin (TPO) to its receptor MPL on the cell surface. MKs undergo an altered cell cycle, termed endomitosis, in which they skip cytokinesis effectively doubling their DNA content with each mitotic cycle, which results in polyploid cells of up to 64N (*2*). Mature MKs can release long cytoplasmic extensions, called proplatelets, into vessel sinusoids, which further mature into platelets in the circulation (*3*). While it was long believed that the main function of MKs was to produce platelets, seminal studies have since highlighted that MKs are also important regulators of the bone marrow niche by secreting cytokines such as platelet factor 4 (PF4) and transforming growth factor β1 (TGFβ1) (*4, 5*), presumably from their most abundant granule type, α-granules. The notion that several MK subsets with different functions exist was recently supported by single-cell RNA sequencing (scRNA Seq) of murine and human bone marrow-derived MKs, revealing up to five different MK clusters with distinct functions, one of which was termed niche-regulating due to an enhanced expression of extracellular matrix proteins and cytokines including TGFβ1 (*6–8*).

Philadelphia chromosome-negative myeloproliferative neoplasms (MPNs) consist of three entities: polycythemia vera (PV), essential thrombocythemia (ET) or primary myelofibrosis (PMF). Both PV and ET have the potential to transform into secondary myelofibrosis (MF), which significantly increases mortality (*9, 10*). The vast majority of MPNs are caused by somatic mutations within the genes encoding for the TPO receptor *MPL*, its downstream effector Janus kinase 2 (*JAK2*) or the calcium-dependent chaperone protein calreticulin (*CALR*) (*11*). The main hallmarks of MF are progressive bone marrow fibrosis causing pancytopenia and severe splenomegaly due to extramedullary hematopoiesis. The current standard of care for low-risk MF are JAK2 inhibitors (four approved inhibitors, among them ruxolitinib), which can alleviate disease symptoms and increase survival (*12*). However, the effects of JAK2 inhibition are often temporary since patients can become resistant, thus demanding for novel treatment strategies. Of note, most novel treatment strategies are currently targeting dysfunctional MK maturation, such as inhibitors of the cell cycle regulator Aurora kinase A (*13, 14*).

MF is characterized by the accumulation of small, dysmorphic MKs of lower ploidy, which are the major source of the high levels of TGFβ1 observed in the bone marrow of MF patients (*15–17*). A comparison of the MK transcriptome of pre-fibrotic MF versus overt MF patients recently emphasized an increased abundance of MKs enriched in extracellular matrix and cytokines (*18*). The importance of MKs in promoting marrow fibrosis is highlighted in mouse models with defective α-granule biosynthesis (in mice lacking neurobeachin-like 2) (*19*), altered endosome recycling in MKs (in mice lacking the GTPase dynamin2) (*20*), or lack of inhibitory stimuli (upon mutation of the MK and platelet inhibitory receptor *MPIG6B*) (*21, 22*). TGFβ1 is a profibrotic cytokine that is assembled as a precursor in a covalent complex with the signal peptide latency-associated peptide (LAP) as well as other latent TGFβ1 binding proteins that are cleaved after TGFβ1 is secreted into the extracellular space (*23*). Transplantation of bone marrow derived from TGFβ1-deficient mice prevented fibrosis in a TPO overexpression mouse model of MF (*24*), highlighting the essential role of TGFβ1 in promoting fibrosis. Non-canonical TGFβ1 signaling within mesenchymal stromal cells is causative for fibrosis progression in mouse models of MF (*25*). Direct inhibition of TGFβ using a TGFβ1/3-specific protein trap (AVID200) has demonstrated beneficial results in mouse models of MF and is currently being tested in early-phase clinical trials (*26, 27*). While the contribution of MKs to MF progression is widely appreciated, the underlying mechanisms of increased cytokine secretion by neoplastic MKs are unresolved.

Autophagy is a conserved cytoplasmic self-digestion process necessary for the degradation or recycling of organelles or protein aggregates (*28*). Secretory autophagy - the fusion of autophagosomes with the plasma membrane rather than lysosomes - has been demonstrated as an unconventional secretion mechanism for leaderless proteins, best described for the activating cytokine interleukin (IL) 1β (*29, 30*). Importantly, inhibition of autophagy has been demonstrated to reduce the secretion of TGFβ1 from fibroblasts (*31*). Along with the vital autophagy regulator mammalian target of rapamycin, autophagy in fibroblasts is controlled by Rho kinase 1 (ROCK1), an effector kinase of the small GTPase RhoA (*32*). Of note, the RhoA pathway has recently been shown to be upregulated in MKs in a mouse model of MF driven by TPO overexpression (*33*), and RhoA guanine exchange factor (GEF) expression is significantly higher in MKs of MF patients (*18, 34*), suggesting a possible connection between RhoA activity and MF progression.

Here, we demonstrate that TGFβ1 is not, as previously suggested (*35*), stored within MK α-granules, but is rather released via secretory autophagy downstream of the small GTPase RhoA. *In vitro*, inhibition of secretory autophagy either directly or through interfering with RhoA/ROCK signaling upstream of autophagosome formation, reduced TGFβ1 secretion and led to its intracellular accumulation alongside cytosolic autophagy marker light chain 3B-I (LC3B-I). Deletion of either autophagy-related gene 5 (*Atg5)* or *RhoA* ameliorated disease progression by reducing TGFβ1 and IL1β cytokine burden and fibrosis in an MPL^W515L^-driven MF mouse model. Lastly, mono- or combination treatment of the autophagy inhibitor hydroxychloroquine or the ROCK inhibitor Y27632 with a JAK2 inhibitor in MPL^W515L^-transplanted mice significantly reduced TGFβ1 and IL1β levels and collagen deposition in the bone marrow and significantly ameliorated fibrosis. Together, our data revealed a novel secretion mechanism of TGFβ1 and suggests potential therapeutic benefits of targeting autophagy to limit the progression of MF.

### Methods and Materials Mice

C57BL/6N mice (#027C57BL/6) were acquired from Charles River Laboratories (Worcester, United States). BALB/cJ mice were acquired from Jackson Laboratories (#000651). *Tpo^−/-^* and *Mpl^−/-^* mice were kindly provided by Prof. Ann Mullally (Brigham and Women’s Hospital, Boston, Massachusetts). *RhoA^fl/fl^* mice were kindly provided by Prof. Kimberley Tolias (Baylor College of Medicine, Houston, Texas). *Rhoa^fl/fl,PF4Cre^* (referred to as *RhoA^MK-KO^*) mice were generated by crossing *RhoA^fl/fl^* mice with PF4-Cre transgenic mice (Jackson Laboratories, C57BL/6-Tg(PF4-icre)Q3Rsko/J). Myxovirus resistance 1 (Mx1)-Cre-positive, *Atg5^fl/fl^*mice *(Atg5^fl/fl,Mx1Cre^*, referred to as *Atg5^HSC-KO^*) were generated in the lab of Prof. Robert Signer (University of California San Diego, CA). To induce deletion of *Atg5* from hematopoietic stem cells, mice were administered with double stranded RNA [poly(I):poly(C)] at 10 weeks of age and bone marrow was shipped and used for transplantation 4 weeks later. *Atg5^fl/fl,Mx1Cre^*mice were housed in the animal facilities at the University of California San Diego, CA. All other mice were housed in the animal facilities at Boston Children’s Hospital, Boston, MA. All animal work was approved by the International Animal Care and Use Committee at Boston Children’s Hospital, Boston, MA (00001248).

### Genotyping

*RhoA* floxed gene and Cre expression were assessed by PCR in DNA retrieved from ear punches. DNA was lysed using DirectPCR lysis reagent (102-T, Viagen Biotech) and samples were analyzed using a pre-mixed PCR mix (PB10.13-02, PCR Biosystems) and the following primers:

PF4-Cre:

PF4 for: 5’ CCC ATA CAG CAC ACC TTT TG 3’
PF4 rev: 5’ TGC ACA GTC AGC AGG TT 3’

*Rhoa*:

RhoA flox for: 5’ AGG GTT TCT CTG TAC GGT AGT C 3’
RhoA flox rev: 5’ GCA GCT AGT CTA ACC CAC TAC A 3’

### Blood parameters

Mice were retro-orbitally bled into EDTA-coated tubes under isoflurane anesthesia using heparinized capillaries. Blood cell counts were assayed at an automated blood cell analyzer (Sysmex Corporation).

### Isolation of bone marrow megakaryocytes

Mice were sacrificed by CO_2_ asphyxiation followed by cervical dislocation, and femora, tibiae and iliac crests were isolated. Cells were isolated by centrifugation for 40 seconds at 2500 x g as previously described (*36*). Primary MKs were isolated as described before (*37*). Briefly, red blood cells were lysed in ACK buffer (A1049201, Gibco) and cells were passed through a 70 µm cell strainer. Afterwards, the cell suspension was consecutively filtered using 20 and 15 µm cell strainers (43-50020-03 and 43-50015-03, pluriSelect) and megakaryocytes were retrieved by inverting strainers. MKs were immediately fixed in 4% PFA/PBS containing 0.1% Tween20 and stored in PBS until further processing.

### Isolation and culture of murine bone marrow-derived megakaryocytes

Bone marrow cells were isolated from C57BL/6N mice by centrifugation as described above (*36*). Cells were passed through a 70 µm cell strainer and with a rat anti-mouse lineage panel (133307, BioLegend), followed by magnetic bead isolation using Dynabeads^TM^ (11415D, ThermoScientific). Lineage-depleted hematopoietic stem and progenitor cells (HSPCs) were cultured in DMEM containing 50 ng mL^−1^ TPO. For proplatelet formation, 100 anti-thrombin units (ATU) of recombinant hirudin (ARE120A, Aniara Diagnostic) were added to the cells. For inhibitor studies, MKs were treated with 5 µM CCG1423 (HY-13991), 500 nM Y27632 (HY-10583), 50 µM endosidin2 (HY-120821), 5 µM Verteporfin (HY-B0146) or 5 µM Hydroxychloroquine (HY-B1370, all MedChem Express). After 72 hours, cells were spun down at 300 x g for 5 min, supernatant was frozen down at -20°C and cells were either used for flow cytometry or mature MKs were enriched using size exclusion filtration.

### Retrieval of bone marrow fluid

For retrieval of bone marrow fluid, mice were sacrificed, and femurs, tibias and iliac crests were isolated. Bones were spun out into 100 µL PBS, bone marrow fluid was retrieved, centrifuged at 10,000 x g for 8 min and frozen at -20°C.

### Analysis of hematopoietic stem and progenitor cells by flow cytometry

Mice were sacrificed by CO_2_ asphyxiation followed by cervical dislocation, and femora, tibiae and iliac crests were isolated. Cells were isolated by centrifugation for 40 seconds at 2500 x g as previously described (*36*). Red blood cells were lysed in ACK buffer (A1049201, Gibco) and cells were passed through a 100 µm cell strainer. Cells were counted using an automated cell counter (Countess 3, ThermoFisher Scientific). Cells were stained for a lineage panel (CD3 (155612, BioLegend), Ly-6G (127612, BioLegend), CD11b (101224, BioLegend), B220 (103227, BioLegend), Ter-119 (116232, BioLegend), Nk1.1 (108722, BioLegend), CD19 (152416, BioLegend) c-kit (135138, BioLegend), Sca-1 (108134, BioLegend), CD150 (115916, BioLegend), CD48 (103432, BioLegend), CD135 (135306, BioLegend), CD41 (133926, BioLegend) and CD42d (148506, BioLegend). Live cells were distinguished using Zombie Aqua (423102, BioLegend). HSPC populations were identified by spectral flow cytometry (Cytek Aurora) as previously described (*38, 39*).

### Cryosectioning and immunofluorescence staining

Femurs were isolated as described above and fixed using 4% PFA/PBS overnight. Afterwards, femora were moved into 10% sucrose/PBS (24h at 4°C), followed by 20% and 30% sucrose. Dehydrated femurs were embedded in a water-soluble embedding medium and frozen down at -20°C. Using a tape transfer system (*40*), 10 µm sections were retrieved at a cryostat (Leica Biosystems). Sections were rehydrated in PBS for 20 min, non-specific binding was blocked using 10% donkey serum and sections were stained for either CD105 (AF1320, R&D Systems), laminin (L9393, Sigma-Aldrich), LAP (AF-246-NA, R&D Systems), von Willebrand factor (vWF) (PA5-16634, ThermoFisher Scientific), collagen I (ab21286, abcam), collagen IV (ab6586, abcam), CD41 (133902, BioLegend) or CD42b (M-051-0, Emfret Analytics). Nuclei were stained using 4′,6-diamidino-2-phenylindole (DAPI). Slides were washed in PBS containing 0.1% TritonX100 and mounted using Fluoroshield mounting media (F6182, Sigma-Aldrich). Sections were imaged at a Zeiss LSM880 confocal microscope (20x objective). Whole femora were imaged and MKs were quantified using an automated imaging platform and Gen5 software (Lionheart, Biotek; 4x objective). Mean fluorescent intensities (MFIs) and collagen deposition were quantified using Fiji Software (Version 2.14.0, NIH).

### Immunofluorescence staining of megakaryocytes and platelets

Primary bone marrow-derived or *in vitro*-cultured MKs were isolated as described above and fixed in 4% PFA/PBS containing 0.1% Tween20 for 20 min at RT. Cells were spun down at 300 x g for 5 min and nonspecific antibody binding was blocked using 3% BSA/PBS. Proplatelet-forming cells were cultured in 8-well chambers (155360, Nunc Lab-Tek), coated with an anti-mouse-CD31 antibody (102502, BioLegend) for 24h and fixed as described above. For platelet isolation, mice were anaesthetized using isoflurane and bled into EDTA-coated tubes (365974, BD). Modified Tyrode’s Buffer (137 mM NaCl, 2.7 mM KCl, 1.0 mM MgCl2, 0.18 mM CaCl2, 20 mM HEPES, and 5.6 mM glucose, pH 7.4) was added and blood was centrifuged for 8 min at 100 x g. The platelet fraction was retrieved, washed in Tyrode’s Buffer and centrifuged at 100 x g again. The platelet fraction was removed and centrifuged at 750 x g for 5 min. Platelet pellets were resuspended in Tyrode’s buffer containing 0.2 µg mL^−1^ PGE_1_ (P5515, Sigma-Aldrich) and counted by flow cytometry. Coverslips were coated overnight at 4°C with 100 µg mL^−1^ fibrinogen and platelets were allowed to adhere for 15 min. MKs and platelets were stained overnight using antibodies against LC3B (PA5-35195, Invitrogen), LAP (AF-246-NA, R&D Systems), vWF (PA5-16634, ThermoFisher Scientific), proplatelet basic protein (PPBP) (PA5-47947, Invitrogen), PF4 (AF595, Novus Biologicals), α-tubulin (322588, ThermoFisher Scientific) and CD42b (M-051-0, Emfret Analytics), nuclei were stained using DAPI. Cells were imaged at a Zeiss LSM880 confocal microscope (40x or 63x objective). Mean fluorescent intensities (MFIs) were quantified using Fiji Software (Version 2.14.0, NIH).

### Analysis of megakaryocyte maturation and ploidy by flow cytometry

Lineage-depleted, bone marrow-derived cells were cultured in TPO-containing media for 4 days. Cells were centrifuged at 300 x g, resuspended in MACS buffer (Miltenyi) and stained for CD41 (133904, BioLegend) and CD42d (148504, BioLegend) on ice for 30 min. The number of double-positive cells was assessed by flow cytometry (Accuri C6 plus, BD Biosciences). For ploidy analysis, HSPCs were cultured in the presence of Dimethyl sulfoxide (DMSO) or Y27632 as described above. After 72 hours, cells were fixed and permeabilized in 70% ethanol for 30 min on ice. Cells were treated with RNAse A (EN0531, ThermoFisher Scientific), stained with anti-CD41-FITC antibodies (133904, BioLegend) and propidium iodide (P1304-MP, Sigma-Aldrich) for 30 min on ice, and ploidy distribution was assessed using flow cytometry (BD Accuri C6 plus).

### Enzyme-linked immunosorbent assays

Cytokine levels in bone marrow fluid or cell culture supernatants were determined using a mouse TGF-beta 1 DuoSet ELISA Kit (DY1679, R&D Systems), a mouse CXCL4/PF4 DuoSet ELISA Kit (DY595, R&D Systems) or a mouse IL-1beta DuoSet ELISA Kit (DY401, R&D Systems).

### MPL^W515L^ virus production

The MSCV-IRES-EGFP-MPL^W515L^ plasmid was kindly provided by Dr. Ross Levine (Memorial Sloan Kettering Cancer Center, New York) (*41*). For virus production, human embryonic kidney cells (HEK 293T) cells were transfected with 15 µg of the MSCV-MPL^W515L^ plasmid or an MSCV-EGFP plasmid and 15 µg of a pCL-eco packaging plasmid (https://www.addgene.org/12371/) using 50 µL lipofectamine 2000® (11668027, Invitrogen). Cells were plated at a density of 5×10^6^ cells in 10 cm dishes, and the media was changed 24h after plating. Cells were incubated for another 48-72h. Viral supernatant was collected, and cellular debris was removed by centrifugation at 1000 x g for 5 min. Viral supernatant was aliquoted and stored at -80°C until further use.

### MPL^W515L^-transplant model of myelofibrosis

MPL^W515L^-transplant model was performed as previously described (*41*). Briefly, bone marrow cells were retrieved from donor mice by crushing femurs, tibiae, iliac crests, humeri, sterna and spines. Cells were passed through a 100 µm cell strainer and centrifuged at 300 x g for 5 min. Red blood cells were lysed in ACK buffer (A1049201, Gibco), cells were passed through a 40 µm cell strainer, and centrifuged at 300 x g for 5 min. CD117^+^ cells were enriched using CD117 microbeads (130-091-224, Miltenyi Biotec) as per the manufacturers’ instructions and cultured at 1×10^6^ cells per well overnight at 37°C with 5% CO_2_ in stem cell media (MEM-α supplemented with 10% FBS, 100 U penicillin and streptomycin, 10 ng/mL murine TPO, 10 ng/mL murine IL-3, 10 ng/mL murine FLT3L, and 100 ng/mL murine SCF). The following day, 5×10^6^ cells per tube were spun down at 300 x g for 5 min and resuspended in 1 ml of MSCV-MPL^W515L^-IRES-EGFP (or MSCV-EGFP) virus containing polybrene (TR-1003-G, Sigma-Aldrich) and centrifuged at 1200 x g and 30°C for 90 min. Cells were incubated for another 90 min in the incubator at 37°C, spun down and resuspended in MEM-α with 10% FBS, 100 U penicillin and streptomycin, 10 ng/mL murine TPO, 10 ng/mL murine IL-3, 10 ng/mL murine FLT3L, and 100 ng/mL murine SCF. Cells were used for transplantation the following day. 1×10^6^ cells were injected into lethally irradiated recipient mice (2 x 450 rad for BALB/CJ mice; 2 x 550 rad for C57BL/6J mice, four hours apart). Mice were kept on prophylactic antibiotics (trimethoprim-sulfamethoxazole) for two weeks after transplantation. For pharmacological studies, BALB/cJ mice were given 30 mg/kg Y27632 (HY-10583, MedChem Express), 50 mg/kg Hydroxychloroquine (HY-B1370, MedChem Express) and/or 60 mg/kg Ruxolitinib (HY-50856) once daily by oral gavage for 2-3 weeks depending on disease severity. Mice were closely monitored and sacrificed when they became moribund (3-4 weeks after transplantation), or when 90% of platelets (CD41^+^) were EGFP^+^ as assessed by flow cytometry.

### Immunoblotting

Megakaryocytes were cultured *in vitro* for four days as described above and lysed using 1x Radio-Immunoprecipitation Assay (RIPA) buffer. Protein concentrations were assessed using the Pierce BCA Protein Assay Kit (ThermoFisher Scientific). 15 µg of protein were loaded onto Sodium dodecyl-sulfate polyacrylamide gels (SDS-PAGE). For platelet immunoblots, 800,000 platelets µL^−1^ were lysed in 1x RIPA buffer and SDS-PAGE and turbo transfer were performed (Bio-rad Laboratories Inc). Polyvinylidene fluoride (PVDF) membranes were probed for LC3B (PA5-35195, Invitrogen), RhoA (sc-418, Santa Cruz Biotechnology Inc) and GAPDH (5174S, Cell Signaling Technologies).

### Statistics

All results are displayed as mean ± *standard deviation* (SD). The distribution of data was assessed with a Shapiro-Wilk test. Differences were statistically analyzed as stated in the figure legends using either unpaired, two-tailed Student’s t-tests, one-or two-way ANOVA with Dunnett’s testing, Sidak’s or Tukey correction for multiple comparisons. P*-*values <0.05 were considered statistically significant.

## Results

### TGFβ1 localizes to MK and platelet autophagosomes

Among other cytokines, MK-derived TGFβ1 has been implicated as a major regulator of HSC quiescence, which prompted us to validate these findings in mice with reduced MK and platelet counts. As a model for this, we used mice lacking either TPO (*Tpo^−/-^*) or its receptor MPL (*Mpl^−/-^*), both of which present with an approximately 90% reduction in MK and platelet counts (*42, 43*). Both models exhibited markedly reduced levels of TGFβ1 in their bone marrow fluid (**Figure 1A**) validating that MKs contribute to homeostatic levels of extracellular TGFβ1 in the bone marrow. Recent advances in single-cell omics have identified subsets of MKs with niche-regulating capacities exhibiting lower ploidy and smaller cell size (*6, 7*). To investigate whether TGFβ1 levels in MKs correlated with maturation, we stained femoral cryosections of wild-type (*WT*) mice for the mature MK marker CD42b, the TGFβ1-associated protein LAP and the prominent α-granule protein vWF (**Figure 1B**). While vWF expression levels positively correlated with CD42b (**Figure 1C**), in line with an increase in α-granule biogenesis observed throughout MK maturation (*19, 44*), we detected a negative correlation between LAP and CD42b staining intensity (**Figure 1D**). To assess cytokine release *in vitro*, we cultured HSPCs in the presence or absence of TPO and assessed TGFβ1 and PF4 levels after 72 hours of culture. While there was limited release of TGFβ1 from MKs into the supernatant (**Figure 1E**), PF4 was largely absent under no-TPO conditions and increased significantly in the presence of MKs (**Figure 1F**), suggesting differential cytokine release kinetics *in vitro*.

**Figure 1.**
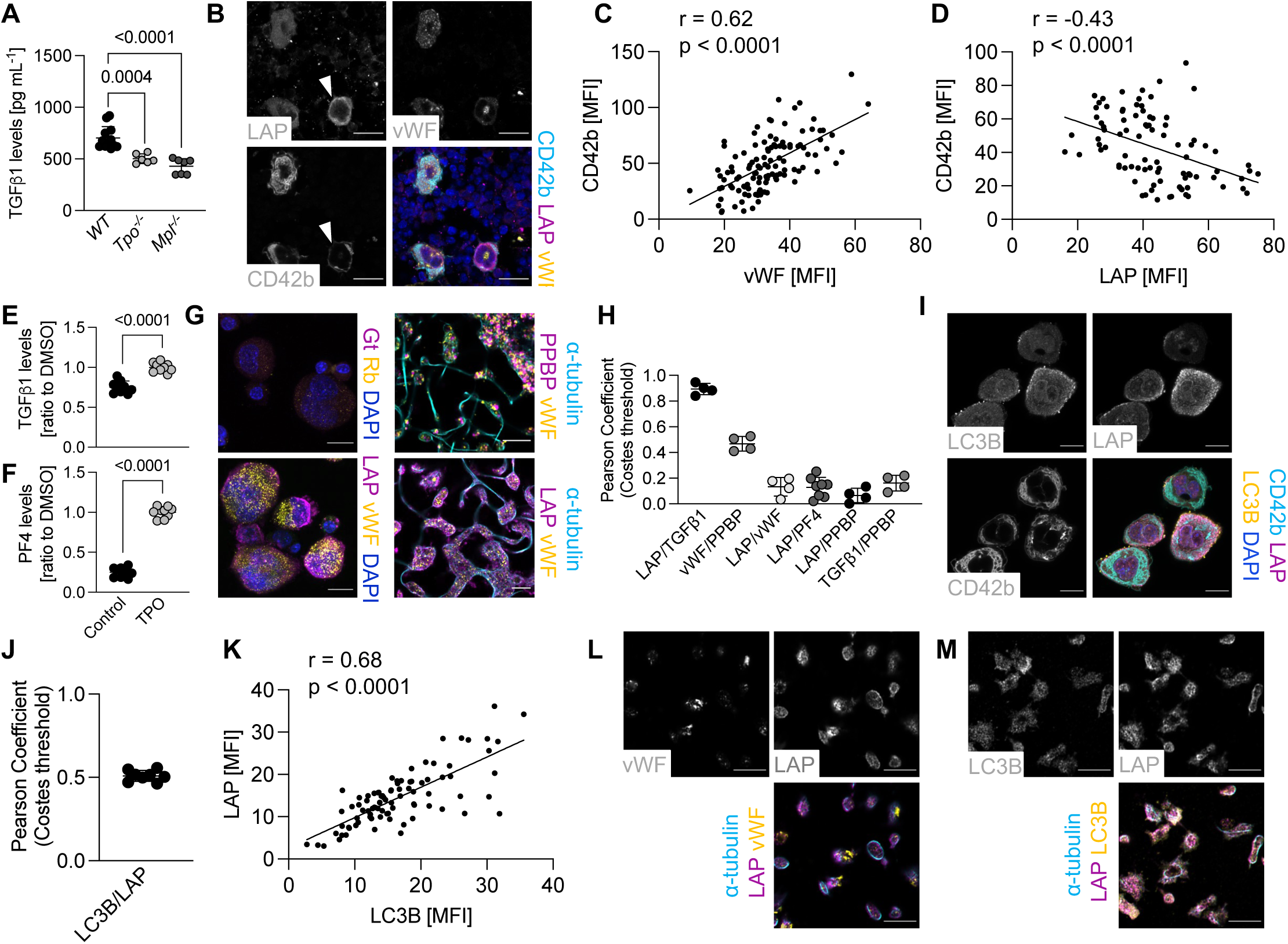
Expression of TGFβ1 is independent of classical α-granule proteins in megakaryocytes. **(A)** Analysis of transforming growth factor β1 (TGFβ1) levels in bone marrow fluid derived from C57BL/6N, thrombopoietin (TPO)-deficient (*Tpo^−/-^*) or MPL-deficient mice (*Mpl^−/-^*). n=6-10 mice. One-way ANOVA with Dunnett’s test for multiple comparisons. **(B)** Visualization of latency-associated peptide (LAP)/TGFβ1, von Willebrand Factor (vWF) and the megakaryocyte (MK) marker CD42b in femoral cryosections. Nuclei were counterstained using DAPI. Scale bars: 20 µm. **(C, D)** Correlation analysis between CD42b and vWF **(C)** and CD42b and LAP/TGFβ1 **(D)**. Each dot represents one MK. n=2 mice. **(E, F)** Analysis of TGFβ1 and PF4 levels in cell culture supernatant after incubation of HSPCs in the absence (Control) or presence of TPO. Values were normalized to TPO values. n=8 mice. Unpaired, two-tailed Student’s t-test. **(G)** Visualization of LAP/TGFβ1, vWF and proplatelet basic protein (PPBP) in round and proplatelet-forming MKs. Goat (gt) and rabbit (rb) isotype antibodies served as staining controls. Nuclei were counterstained using DAPI. Scale bars: left: 10 µm; right: 5 µm. **(H)** Pearson colocalization coefficient analysis using Costes threshold of proplatelet-forming MKs imaged using super-resolution microscopy (Airyscan Leica LSM880, 63x objective). Each dot represents one MK. LAP/TGFβ1 served as a positive control. **(I)** Visualization of LC3B, LAP/TGFβ1 and CD42b in native bone marrow MKs. Nuclei were counterstained using DAPI. Scale bars: 15 µm. **(J)** Pearson colocalization coefficient analysis using Costes threshold for LC3B and LAP/TGFβ1. Each dot represents one MK. n=2 mice. **(K)** Correlation analysis between LAP/TGFβ1 and LC3B in cultured, bone marrow-derived MKs. Each dot represents one MK. n=3 mice. **(L, M)** Visualization of vWF and LAP/TGFβ1 **(L)** as well as LC3B and LAP/TGFβ1 **(M)** in platelets adhered on a fibrinogen-coated surface. Platelets were stained for α-tubulin to highlight the cytoskeleton. Scale bars: 5 µm. All data are presented as mean ± SD.

α-granules are the most abundant granule type in MKs and contain around 300 molecules, among them TGFβ1 as previously suggested (*35*). However, since we observed differential cytokine release kinetics for TGFβ1 and PF4, we hypothesized that TGFβ1 might be stored independently of classical α-granules. To test this, we performed super-resolution confocal microscopy on round and proplatelet-forming MKs (**Figure 1G**). When visualizing granular content, we observed no colocalization of LAP with either of the tested α-granule proteins (PF4, vWF or proplatelet basic protein (PPBP)) as assessed using a Pearson Costes colocalization coefficient (**Figure 1H**). The two different α-granule proteins vWF and PPBP on the other hand co-localized to a much higher extent, suggesting they are present within the same organelle, as did LAP and TGFβ1, using two different antibodies. It has previously been demonstrated that TGFβ1 is released via secretory autophagy in fibroblasts (*31*). To therefore test whether TGFβ1 localized to autophagosomes, we isolated native MKs from the bone marrow and visualized LAP and the autophagy marker LC3B (**Figure 1I**). As assessed using Pearson Costes colocalization, we observed a high colocalization of LAP and LC3B (**Figure 1J**), suggesting that the cytokine is at least partially localized to autophagosomes in mature MKs. When analyzing cultured MKs, we observed a significant positive correlation between LAP and LC3B expression (**Figure 1K**), implying overlapping expression patterns. To verify our findings, we isolated platelets from *WT* mice, let them adhere to a fibrinogen-coated surface and visualized vWF and LAP (**Figure 1L**) or LC3B and LAP (**Figure 1M**). Similar to our observations in MKs, TGFβ1/LAP was expressed distinctly from conventional α-granules, and instead exhibited substantial colocalization with LC3B. In summary, these findings raise the possibility that TGFβ1 is stored independently of classical α-granules in MKs and may be secreted via an unconventional pathway.

### TGFβ1 secretion is dependent on secretory autophagy downstream of RhoA/Rho kinases

Autophagosomes originate from subdomains of the endoplasmic reticulum, referred to as omegasomes (*45*) and their formation and maturation can be targeted using a variety of inhibitors. The benzoporphyrin derivative Verteporfin (VP) inhibits early autophagosome formation (*46*), while the antimalarial drug hydroxychloroquine (HQ) inhibits the fusion of autophagosomes with lysosomes (**Figure 2A**) (*47*). To test the effect of autophagy inhibition on MKs *in vitro*, we treated HSPCs with the two autophagy inhibitors and assessed MK maturation and TGFβ1 levels in the supernatant after culturing the cells for 72 hours in the presence of TPO. Previous studies suggested that autophagy inhibition impaired MK maturation *in vitro* (*48, 49*). In line with this, autophagy inhibition using HQ and VP reduced the percentage of CD41 and CD42d double-positive cells (**Figure 2B**), confirming the important role of autophagy during megakaryopoiesis. To test whether inhibition of autophagy reduced the release of TGFβ1, we next assessed TGFβ1 levels in the supernatant of treated cells and observed significantly reduced concentrations of TGFβ1 for both tested inhibitors (**Figure 2C**). Of note, when comparing TGFβ1 levels upon HQ or VP treatment to supernatant derived from cells cultured in the absence of TPO they were similar, suggesting that autophagy inhibition diminished TGFβ1 secretion from MKs. Another cytokine that was previously shown to be released via secretory autophagy and is expressed in MKs is IL1β (*29, 30, 50*). We therefore also assessed levels of IL1β in the cell culture supernatant after treatment of HSPCs with autophagy inhibitors and observed reduced levels following HQ and VP treatment (**Figure 2D**). To exclude that the observed effects were due to a general reduction in the amount of mature MKs and overall cytokine secretion, we also assessed levels of the α-granule protein PF4 in the supernatant of treated cells. However, we did not observe differences in PF4 levels after treatment with HQ, and only a mild reduction after VP treatment (**Figure 2E**), suggesting autophagy inhibition selectively affected the release of TGFβ1 and related cytokines.

**Figure 2.**
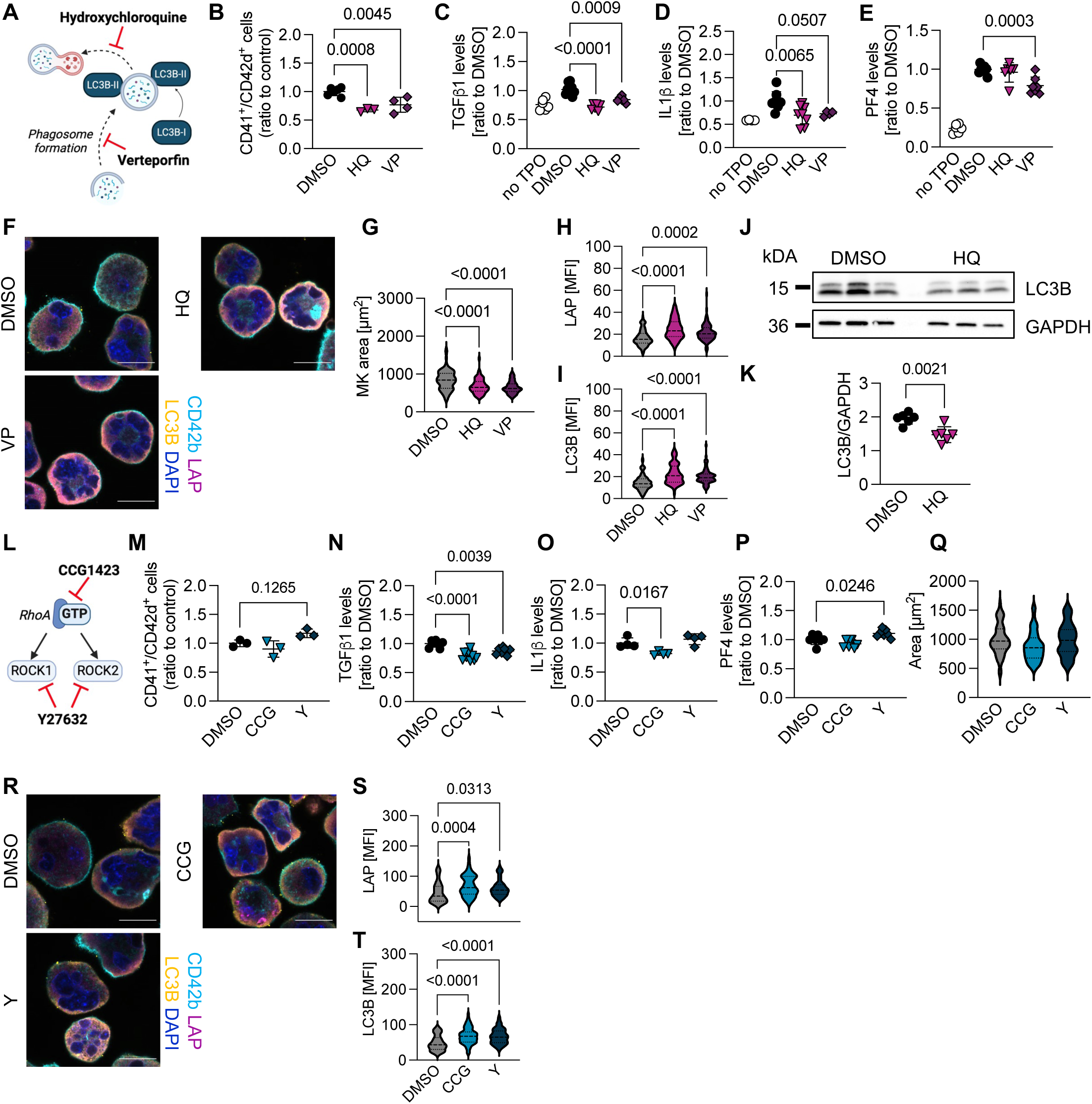
Autophagy or RhoA/ROCK inhibition impairs TGFβ1 and IL1β secretion from megakaryocytes *in vitro*. **(A)** Schematic visualizing how hydroxychloroquine (HQ) and verteporfin (VP) interfere with autophagy. **(B)** Percentage of CD41^+^/CD42d^+^-positive cells after treatment of HSPCs with DMSO, 5 µM HQ or 5 µM VP for 72h was assessed using flow cytometry. Values were normalized to the DMSO control. n=3-4 mice. One-way ANOVA with Dunnett’s test for multiple comparisons. **(C-E)** Analysis of TGFβ1 **(C)**, IL1β **(D)** and PF4 levels **(E)** in cell culture supernatant after treatment of HSPCs with DMSO, 5 µM HQ or 5 µM VP for 72h. Supernatant derived from cells cultured in the absence of TPO were included as a control. Values were normalized to DMSO values. n=4-6 mice. One-way ANOVA with Dunnett’s test for multiple comparisons. **(F)** Visualization of LC3B, LAP/TGFβ1 and CD42b in enriched bone marrow-derived MKs treated with DMSO, HQ or VP for 72h. Nuclei were counterstained using DAPI. Scale bars: 20 µm. **(G-I)** MK area **(G)**, LAP/TGFβ1 **(H)** and LC3B MFI **(I)** were analyzed in cultured MKs. At least 70 MKs were analyzed per condition. n=3. One-way ANOVA with Dunnett’s test for multiple comparisons. **(J, K)** Immunoblot **(J)** and densitometric analysis **(K)** of enriched bone marrow-derived MKs treated with DMSO or HQ for 72h. n=6. Unpaired, two-tailed Student’s t-test. **(L)** Schematic on the inhibitory function of CCG1423 (CCG) and Y27632 (Y) on RhoA and Rho kinases (ROCKs). **(M)** Percentage of CD41^+^/CD42d^+^-positive cells after treatment of HSPCs with DMSO, 5 µM CCG or 500 nM Y for 72h was assessed using flow cytometry. Values were normalized to the DMSO control. n=3 mice. One-way ANOVA with Dunnett’s test for multiple comparisons. **(N-P)** Analysis of TGFβ1 **(N)**, IL1β **(O)** and PF4 levels **(P)** in cell culture supernatant after treatment of HSPCs with DMSO, CCG or Y for 72h. Supernatant derived from cells cultured in the absence of TPO were included as a control. Values were normalized to DMSO values. n=3-6 mice. One-way ANOVA with Dunnett’s test for multiple comparisons. **(Q, R)** MK area was assessed **(Q)** and LC3B, LAP/TGFβ1 and CD42b were visualized **(R)** in enriched bone marrow-derived MKs treated with DMSO, CCG or Y for 72h. Nuclei were counterstained using DAPI. Scale bars: 20 µm. **(S, T)**, LAP/TGFβ1 **(S)** and LC3B MFI **(T)** were analyzed in cultured MKs. At least 70 MKs were analyzed per condition. n=3. One-way ANOVA with Dunnett’s test for multiple comparisons. Schematics were generated using Biorender.com.

To further verify our results and test whether autophagy inhibition induced intracellular retention of TGFβ1, we visualized CD42b, LAP and the autophagy marker LC3B in enriched MKs after 4 days of treatment with HQ or VP (**Figure 2F**). In line with the decrease in maturation, treated cells exhibited a reduced cell size (**Figure 2G**). We observed an increase in mean fluorescence intensity (MFI) for both LAP and diffuse, cytosolic LC3B (**Figure 2H, I**), suggesting that HQ and VP treatment induced TGFβ1 retention *in vitro*. LC3B exists in two different isoforms, the cytosolic LC3B-I and the membrane-bound LC3B-II, which becomes lipidated upon incorporation into the autophagosome membrane (*51*). To distinguish between both isoforms, we also performed immunoblots of cultured MKs and found reduced levels of LC3B upon treatment with HQ (**Figure 2J, K**). Overall, our data suggest that TGFβ1 secretion by MKs is dependent on autophagic activity.

Previous studies have identified an important role of the small GTPase RhoA and its downstream effector Rho Kinase 1 (ROCK1) in regulating autophagosome maturation (*32, 52*). To investigate the role of RhoA/ROCK signaling in secretory autophagy, we first assessed MK maturation upon treatment of HSPCs with the RhoA inhibitor CCG1423 (CCG) and the pan-ROCK inhibitor Y27632 (Y) (**Figure 2L**). While inhibition of RhoA did not affect MK maturation, assessed as the percentage of CD41/CD42d-positive cells, Y enhanced MK maturation (**Figure 2M**), in line with previous publications (*53*). This was also reflected by an increase in 32n, high-ploidy MKs (**Supplemental Figure 1A**). Importantly, TGFβ1 levels in the supernatant were significantly lower upon inhibition of both RhoA and ROCK (**Figure 2N**). Similarly, IL1β levels were reduced upon RhoA inhibition using CCG (**Figure 2O**). In contrast to TGFβ1 and IL1β, PF4 levels in the supernatant of treated MKs were unaltered upon RhoA and even slightly increased upon ROCK inhibition, in line with the enhanced maturation (**Figure 2P**). We also did not observe differences in MK area (**Figure 2Q**).

To further explore these findings, we visualized cytosolic LC3B and LAP in treated MKs (**Figure 2R**). In line with the previous findings from fibroblasts, both RhoA and ROCK inhibition led to an intracellular accumulation of TGFβ1/LAP and cytosolic LC3B (**Figure 2S, T**). To exclude a general effect on granule content, we also assessed expression of the α-granule protein vWF upon treatment of MKs with CCG and HQ (**Supplemental Figure 1B**) and did not observe accumulation upon either direct autophagy or RhoA inhibition (**Supplemental Figure 1C**). To validate our findings, we also treated HSPCs with the general exocytosis inhibitor endosidin2 that should limit both secretory autophagy as well as conventional granule release (*54*). As expected, exocytosis inhibition reduced both TGFβ1 and PF4 levels in the supernatant (**Supplemental Figure 1D, E**) and led to an intracellular accumulation of both TGFβ1/LAP as well as vWF (**Supplemental Figure 1F-H**). In summary, our findings suggest an autophagy-dependent mechanism of TGFβ1 release by MKs, which is regulated by the small GTPase RhoA and its downstream effector ROCK.

### Increased autophagy and TGFβ1 secretion in a mouse model of MPL^W515L^-induced myelofibrosis

Our *in vitro* data suggested TGFβ1 secretion at least partly depends on secretory autophagy. To test this hypothesis in a mouse model known to exhibit increased TGFβ1 release from MKs, we employed a preclinical model of MF. CD117^+^ HSPC-enriched bone marrow cells were transduced with a retrovirus encoding a mutant version of the TPO receptor MPL (MSCV-MPL^W515L^-IRES-EGFP, referred to as MPL^W515L^), and subsequently transplanted into lethally irradiated mice (*41*). As expected, MPL^W515L^-transplanted mice exhibited markedly higher white blood cell counts compared to mice transplanted with cells transduced with an MSCV-EGFP construct (referred to as Control) (**Figure 3A**), validating that this model recapitulated important disease characteristics representative of a fast-progressing myeloproliferative neoplasm. While platelet counts were still comparable at the time of blood collection (**Figure 3B**), mean platelet volume (MPV) and immature platelet fraction (IPF) were significantly higher in diseased animals (**Figure 3C, D**), suggesting perturbed platelet production. Approximately 90% of platelets in MPL^W515L^-transplanted mice were EGFP-positive three weeks after transplantation, with only 10% in the EGFP control (**Figure 3E**). Similarly, up to 50% of CD45^+^ leukocytes were EGFP-positive in MPL^W515L^-transplanted mice, dependent on disease severity (**Figure 3F**). Moreover, MPL^W515L^-transplanted mice presented with severe splenomegaly (**Figure 3G**) as previously described (*41*). While the expression of the MK marker CD41 was lower in MPL^W515L^-transplanted mice (**Figure 3H, I**), we detected significantly higher levels of TGFβ1/LAP within MKs of diseased animals (**Figure 3J**). MPL^W515L^-transplanted animals further exhibited an accumulation of collagens I and IV in the bone marrow (**Figure 3K-M**). MPL^W515L^-transduced CD117^+^ cells cultured *in vitro* in the presence of TPO also exhibited higher PF4 and TGFβ1 release compared to MSCV-EGFP-transduced controls (**Figure 3N, O**).

**Figure 3.**
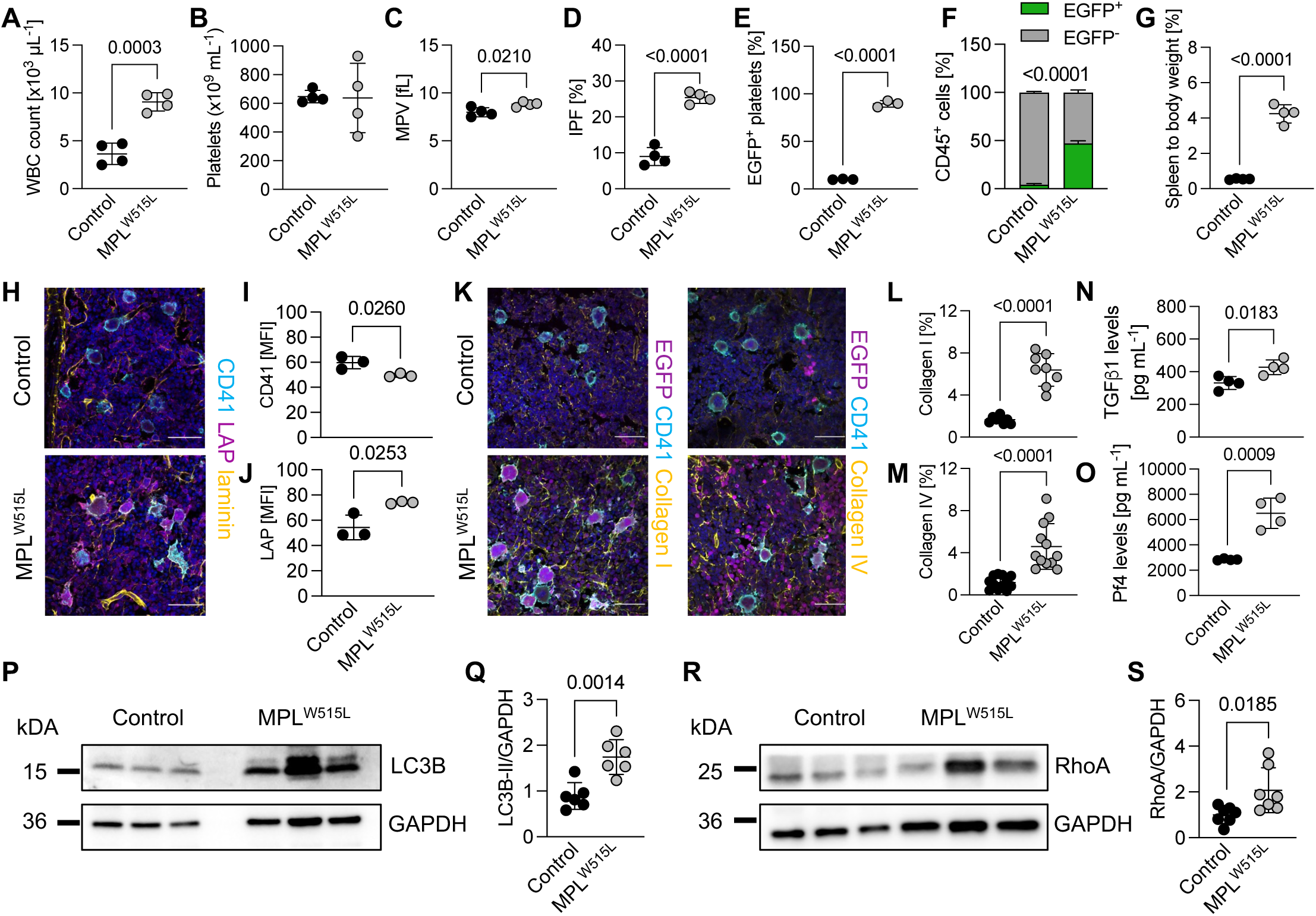
MPL^W515L^ transplant model recapitulates myelofibrosis disease hallmarks and exhibits upregulation of autophagy in the MK lineage. **(A)** White blood cells (WBC) count of mice transplanted with CD117^+^ MSCV-EGFP (Control)- or MSCV-IRES-EGFP-MPL^W515L^ (MPL^W515L^)-transduced cells three weeks after transplantation. n=4 mice. Unpaired, two-tailed Student’s t-test. **(B-D)** Platelet count **(B)**, mean platelet volume (MPV) **(C)** and immature platelet fraction (IPF) **(D)** of mice transplanted with CD117^+^ Control or MPL^W515L^-transduced cells. n=4 mice. Unpaired, two-tailed Student’s t-test. **(E, F)** Percentage of EGFP^+^ platelets **(E)** and CD45^+^ cells **(F)** of mice transplanted with Control or MPL^W515L^-transduced cells three weeks after transplantation. n=4. Unpaired, two-tailed Student’s t-test. **(G)** Spleen to body weight of mice transplanted with Control or MPL^W515L^-transduced cells. n=4 mice. Unpaired, two-tailed Student’s t-test. **(H-J)** Visualization **(H)** and quantification of MFI of CD41 **(I)** and TGFβ1/LAP **(J)** in femoral cryosections of mice transplanted with Control or MPL^W515L^-transduced cells. At least 50 MKs were analyzed per mouse. n=3 mice. Unpaired, two-tailed Student’s t-test. All data are presented as mean ± SD. **(K-M)** Visualization **(K)** and quantification of collagen I **(L)** and collagen IV deposition **(M)** in femoral cryosections of mice transplanted with Control or MPL^W515L^-transduced cells. n=3 mice. Each dot represents one field of view (FOV). Three FOVs were analyzed per mouse. Unpaired, two-tailed Student’s t-test. All data are presented as mean ± SD. **(N, O)** Analysis of TGFβ1 **(N)** and PF4 levels **(O)** in cell culture supernatant of CD117^+^ Control or MPL^W515L^-transduced cells after culture in the presence of TPO for 72h. n=4 mice. Unpaired, two-tailed Student’s t-test. **(P, Q)** LC3B immunoblot **(P)** and densitometric analysis **(Q)** of platelets derived from mice transplanted with CD117^+^ Control or MPL^W515L^-transduced cells. n=6 mice. Unpaired, two-tailed Student’s t-test. **(R, S)** RhoA immunoblot **(R)** and densitometric analysis **(S)** of platelets derived from mice transplanted with CD117^+^ Control or MPL^W515L^-transduced cells. n=6 mice. Unpaired, two-tailed Student’s t-test. All data are presented as mean ± SD.

To test whether autophagy was upregulated in the MK and platelet lineage, we assessed LC3B conversion in platelets from control and MPL^W515L^-transplanted mice by immunoblotting and observed significantly higher levels of lipidated, autophagosome-incorporated LC3B-II in platelets from MPL^W515L^-transplanted mice (**Figure 3P, Q**). In line with recent findings (*33*), we further discovered elevated protein levels of the GTPase RhoA in platelets derived from MPL^W515L^-transplanted mice (**Figure 3R, S**). In summary, we demonstrate that MPL^W515L^-transplanted mice exhibit increased autophagy and recapitulate important disease characteristics to test TGFβ1 secretion mechanisms.

### Deletion of autophagy-related gene 5 from hematopoietic stem cells protects from MPL^W515L^-induced myelofibrosis

Autophagosome formation is mediated by the consecutive activation of a family of proteins referred to autophagy-related genes (ATGs), which are essential for a functional autophagy machinery (**Figure 4A**) (*45*). To investigate the role of secretory autophagy on TGFβ1 secretion from MKs *in vivo*, we utilized mice with an inducible, Mx1-Cre-mediated deletion of the autophagy-regulating protein *Atg5* leading to excision of the gene from HSPCs upon administration of double stranded RNA (*55*). To assess how *Atg5* deletion affected megakaryopoiesis and TGFβ1 secretion at steady state, we first visualized MKs (CD42b), vessels (laminin) and TGFβ1 (LAP) in femoral cryosections of Mx1-Cre-negative (*WT*) and Mx1-Cre-positive (*Atg5^HSC-KO^*) mice (**Supplemental Figure 2A**). While we did not observe differences in either the expression of the late MK marker CD42b or LAP (**Supplemental Figure 2B, C**), MK numbers were slightly higher in *Atg5^HSC-KO^* mice (**Supplemental Figure 2D**). TGFβ1 levels in the bone marrow, however, were unaltered (**Supplemental Figure 2E**), suggesting that autophagy was dispensable for TGFβ1 secretion at steady state.

**Figure 4.**
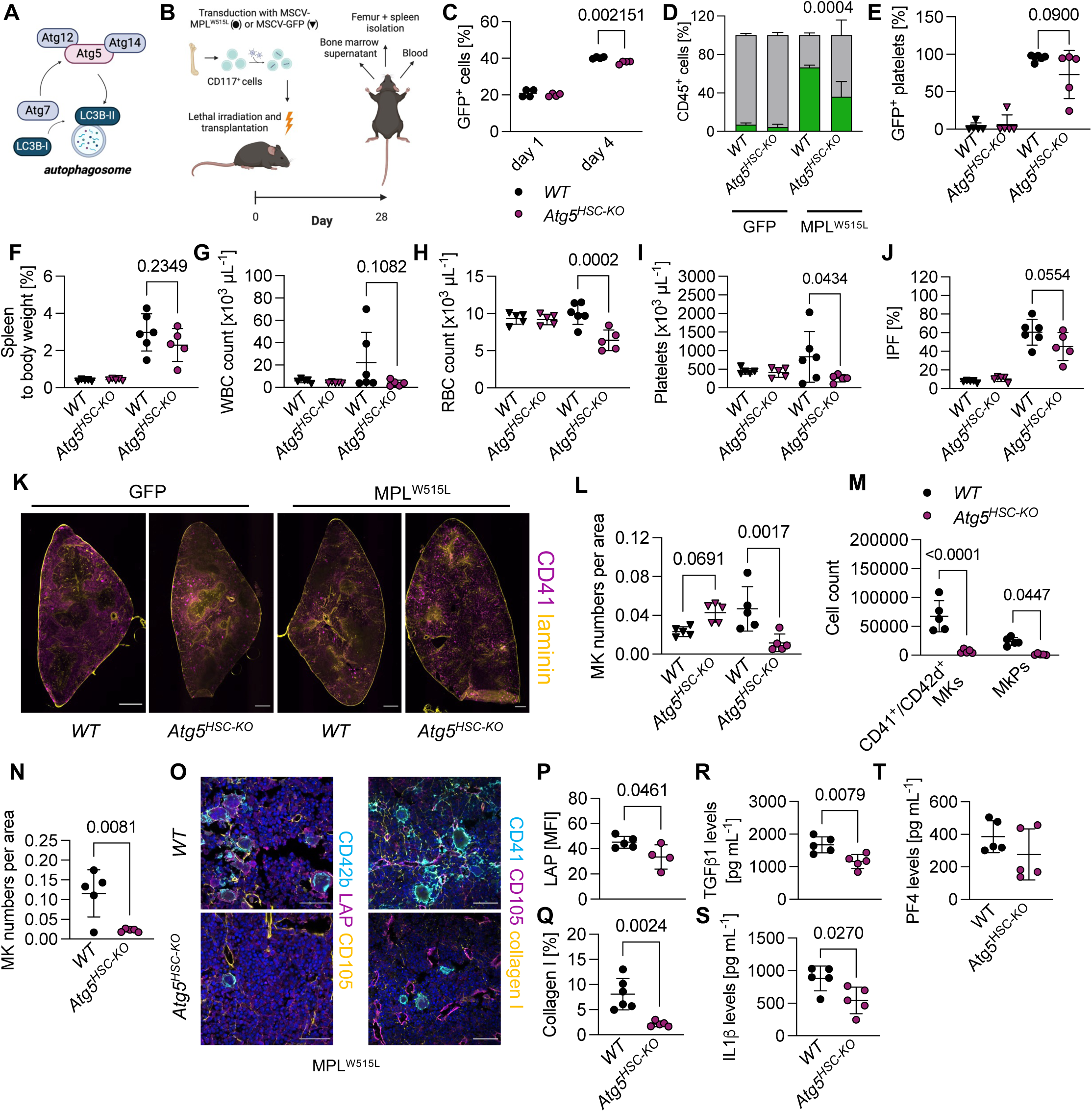
Autophagy deficiency alleviates disease burden in an MPL^W515L^ transplant model of myelofibrosis. **(A)** Schematic of autophagy-related gene 5 (ATG5) in autophagosome formation. **(B)** Schematic of the MPL^W515L^ transplant model in mice. **(C)** Transduction efficiency of CD117^+^ cells derived from *WT* and *Atg5^HSC-KO^* mice transduced with MPL^W515L^ one or four days after transduction. n=4. Multiple, unpaired Student’s t-tests. **(D, E)** Percentage of EGFP^+^ CD45^+^ cells **(D)** and platelets **(E)** of *WT* and *Atg5^HSC-KO^* mice transplanted with Control or MPL^W515L^-transduced cells four weeks after transplantation. n=5-6 mice. Two-way ANOVA with Sidak’s correction for multiple comparisons. **(F)** Spleen to body weight of *WT* and *Atg5^HSC-KO^* mice transplanted with Control or MPL^W515L^-transduced cells. n=5-6 mice. One-way ANOVA with Sidak’s correction for multiple comparisons. **(G, H)** White blood cell (WBC) **(G)** and red blood cell (RBC) counts **(H)** of *WT* and *Atg5^HSC-KO^* mice. n=5-6 mice. One-way ANOVA with Sidak’s correction for multiple comparisons. **(I, J)** Platelet counts **(I)** and immature platelet fraction (IPF) **(J)** of *WT* and *Atg5^HSC-KO^* mice. n=5-6 mice. One-way ANOVA with Sidak’s correction for multiple comparisons. **(K, L)** Visualization **(K)** and quantification **(L)** of splenic MKs of transplanted *WT* and *Atg5^HSC-KO^*mice. n=5-6 mice. One-way ANOVA with Sidak’s correction for multiple comparisons. Scale bars: 500 µm. **(M)** Quantification of MKs and megakaryocyte progenitors (MkP) in the bone marrow of MPL^W515L^ mice transplanted with *WT* or *Atg5^HSC-KO^* cells by flow cytometry. n=5-6 mice. Two-way ANOVA with Sidak’s correction for multiple comparisons. **(N)** Quantification of MK numbers in femoral cryosections of MPL^W515L^ mice transplanted with *WT* or *Atg5^HSC-KO^* cells. n=5 mice. Unpaired, two-tailed Student’s t-test. **(O-Q)** Visualization **(O)** and quantification of intracellular LAP/TGFβ1 **(P)** and collagen I **(Q)** in MPL^W515L^ mice transplanted with *WT* or *Atg5^HSC-KO^*cells. n=5-6 mice. Unpaired, two-tailed Student’s t-test. Scale bars: 50 µm. **(R-T)** Analysis of TGFβ1 **(R)**, IL1β **(S)** and PF4 levels **(T**) in bone marrow fluid of MPL^W515L^ mice transplanted with *WT* or *Atg5^HSC-KO^* cells. n=5-6 mice. Unpaired, two-tailed Student’s t-test. All data are presented as mean ± SD. Schematics were generated using Biorender.com.

To investigate whether reducing autophagy may affect myelofibrosis progression, we transduced *WT-* and *Atg5^HSC-KO^*-derived HSPCs with MSCV-EGFP or MPL^W515L^, transplanted lethally irradiated recipient mice and analyzed control and diseased animals four weeks later (**Figure 4B**). We validated transduction efficiency of cells transduced with MPL^W515L^ and observed no differences between *WT-* and *Atg5^HSC-KO^*-derived CD117^+^ cells one day after transduction with MPL^W515L^ (**Figure 4C**). Of note, the amount of EGFP-positive *Atg5^HSC-KO^* cells was significantly lower when cells were cultured for 72 hours after transduction, suggesting lower proliferation of transduced cells upon *Atg5* deficiency.

Overall, we detected only marginal expression of EGFP in CD45^+^ cells of mice transplanted with the MSCV-EGFP construct. In contrast, the percentage of EGFP-positivity in MPL^W515L^ mice transplanted with *Atg5^HSC-KO^* cells was lower than in the *WT* controls (**Figure 4D**). In line with this, two out of five mice transplanted with *Atg5^HSC-KO^* cells exhibited a reduced number of EGFP-positive platelets (**Figure 4E**). While we did not observe a significant difference in spleen mass (**Figure 4F**), white blood cell counts were not increased and the elevation in red blood cell counts observed in the MPL^W515L^ transplant model were normalized in *Atg5^HSC-KO^*-transplanted mice (**Figure 4G, H**). Similarly, platelet counts were significantly lower in mice transplanted with MPL^W515L^-transduced *Atg5^HSC-KO^* cells, which was further reflected in a trending decrease in the IPF (**Figure 4I, J**), implying lower platelet production. To assess the effect of *Atg5* deletion on MKs in spleens and bone marrow, we next performed immunofluorescence stainings and observed significantly lower splenic MK numbers only in MPL^W515L^ mice transplanted with *Atg5^HSC-KO^* cells (**Figure 4K, L**), suggesting attenuated disease upon lack of *Atg5*. While MK counts in femurs of mice transplanted with *Atg5^HSC-KO^* cells transduced with MSCV-EGFP were only mildly reduced (**Supplemental Figure 2F, G**), MK progenitors (MkPs) and MK numbers in MPL^W515L^ mice transplanted with *Atg5^HSC-KO^* cells were markedly lower compared to mice transplanted with *WT* cells (**Figure 4M**), which was also evident in femoral cryosections (**Figure 4N**), strongly implying alleviated disease burden.

To investigate the effects of *Atg5* deletion on TGFβ1 secretion and fibrosis progression in the bone marrow, we visualized collagen I and LAP in femoral cryosections of MPL^W515L^-transplanted mice (**Figure 4O**). In line with our previous results, we found markedly reduced levels of intracellular LAP in MKs of *Atg5^HSC-KO^*-transplanted mice (**Figure 4P**), accompanied by reduced collagen I deposition (**Figure 4Q**). These differences were echoed by significantly reduced TGFβ1 levels in the bone marrow fluid of *Atg5^HSC-KO^*-transplanted mice (**Figure 4R**). Similarly, levels of IL1β, which is also released via secretory autophagy (*29, 56*) and known to be associated with worsened disease outcome in MF, were lower in *Atg5^HSC-KO^*-transplanted mice (**Figure 4S**). Levels of the α-granule protein PF4 were unchanged (**Figure 4T**), supporting our hypothesis that secretory autophagy specifically regulates the secretion of TGFβ1 and IL1β. Overall, our data suggest that the beneficial effects of autophagy inhibition extend beyond limiting cytokine secretion to improve disease outcome in MF by also affecting proliferation of mutated clones.

### Deletion of RhoA from MKs alleviates fibrosis in MPL^W515L^-transplanted mice

Signaling downstream of the GTPase RhoA has been proposed to regulate autophagy in fibroblasts (*32*) which prompted us to generate mice selectively *RhoA* from MKs and platelets (*RhoA^MK-KO^*). We verified lack of RhoA protein by western blot of platelets derived from *WT* and *RhoA^MK-KO^* mice (**Supplemental Figure 3A**). Like previously generated mice (*57*), RhoA-deficient mice exhibited reduced platelet counts, alongside an increased MPV (**Supplemental Figure 3B, C**). White and red blood cell counts were unaltered (**Supplemental Figure 3D, E**). In line with previous data, we found increased numbers of MkPs and MKs by flow cytometry (**Supplemental Figure 3F, G**), which we verified in cryosections (**Supplemental Figure 3H, I**). MK size on the contrary was slightly lower in RhoA-deficient mice (**Supplemental Figure 3J**). Interestingly, while the expression of the MK marker CD41 was similar in cryosections of *WT* and *RhoA^MK-KO^* mice, LAP/TGFβ1 expression was markedly lower in *RhoA^MK-KO^* mice (**Supplemental Figure 3K, L**), which was in line with reduced levels of TGFβ1 in the bone marrow (**Supplemental Figure 3M**). This contrasted with PF4 levels, which were similar between *WT* and *RhoA^MK-KO^*mice (**Supplemental Figure 3N**), suggesting that *Rhoa* deficiency selectively affected TGFβ1 secretion from MKs at steady state.

To test this in a disease setting, we employed the MPL^W515L^ transplant model using CD117^+^ cells derived from *WT* and *RhoA^MK-KO^* mice. Transduction efficiencies for MSCV-MPL^W515L^ were comparable between *WT* and *RhoA^MK-KO^*cells (**Figure 5A**). Five weeks after transplantation, we did not observe differences in the amount of EGFP-positive CD45^+^ leukocytes (**Figure 5B**) or platelets (**Figure 5C**) for either EGFP- or MPL^W515L^-transplanted mice, suggesting comparable disease progression. Spleen masses were also similar between *WT*-transplanted and *RhoA^MK-KO^*-transplanted mice (**Figure 5D**). When assessing blood cell parameters in transplanted mice, we observed a trending decrease in white blood cell counts in mice transplanted with MPL^W515L^-transduced *RhoA^MK-KO^* cells (**Figure 5E**). While red blood cell counts were similar between *WT* and *RhoA^MK-KO^* (**Figure 5F**), platelet counts were lower in both EGFP- and MPL^W515L^-transplanted mice (**Figure 5G**), reflecting the reduced platelet counts observed in non-transplanted *RhoA^MK-KO^* mice (*see Supplemental Figure 3B*). When assessing MK counts in the spleen (**Figure 5H**), we observed similar MK numbers between *WT* and *RhoA^MK-KO^* in MPL^W515L^-transplanted mice (**Figure 5I**), which was also reflected in femurs (**Figure 5J**). Of note, MK counts in the femurs of mice transplanted with MSCV-EGFP control cells were also similar between *WT* and *RhoA^MK-KO^* (**Supplemental Figure 3O, P**). Importantly, and despite similar MK counts, visualization of collagen I and TGFβ1/LAP in femurs of MPL^W515L^ mice revealed reduced collagen deposition in mice transplanted with *RhoA^MK-KO^* cells compared to *WT* cells (**Figure 5K, L**), which correlated to lower intracellular TGFβ1/LAP levels (**Figure 5M**). While TGFβ1 levels in the bone marrow were not significantly reduced, IL1β levels were markedly lower in mice transplanted with *RhoA^MK-KO^* cells (**Figure 5N, O**). In contrast, PF4 levels in the bone marrow were comparable between *WT* and *RhoA^MK-KO^*-transplanted mice (**Figure 5P**). Overall, our data suggests MK-derived RhoA at least partly contributed to cytokine secretion at steady state and upon myeloproliferative transformation.

**Figure 5.**
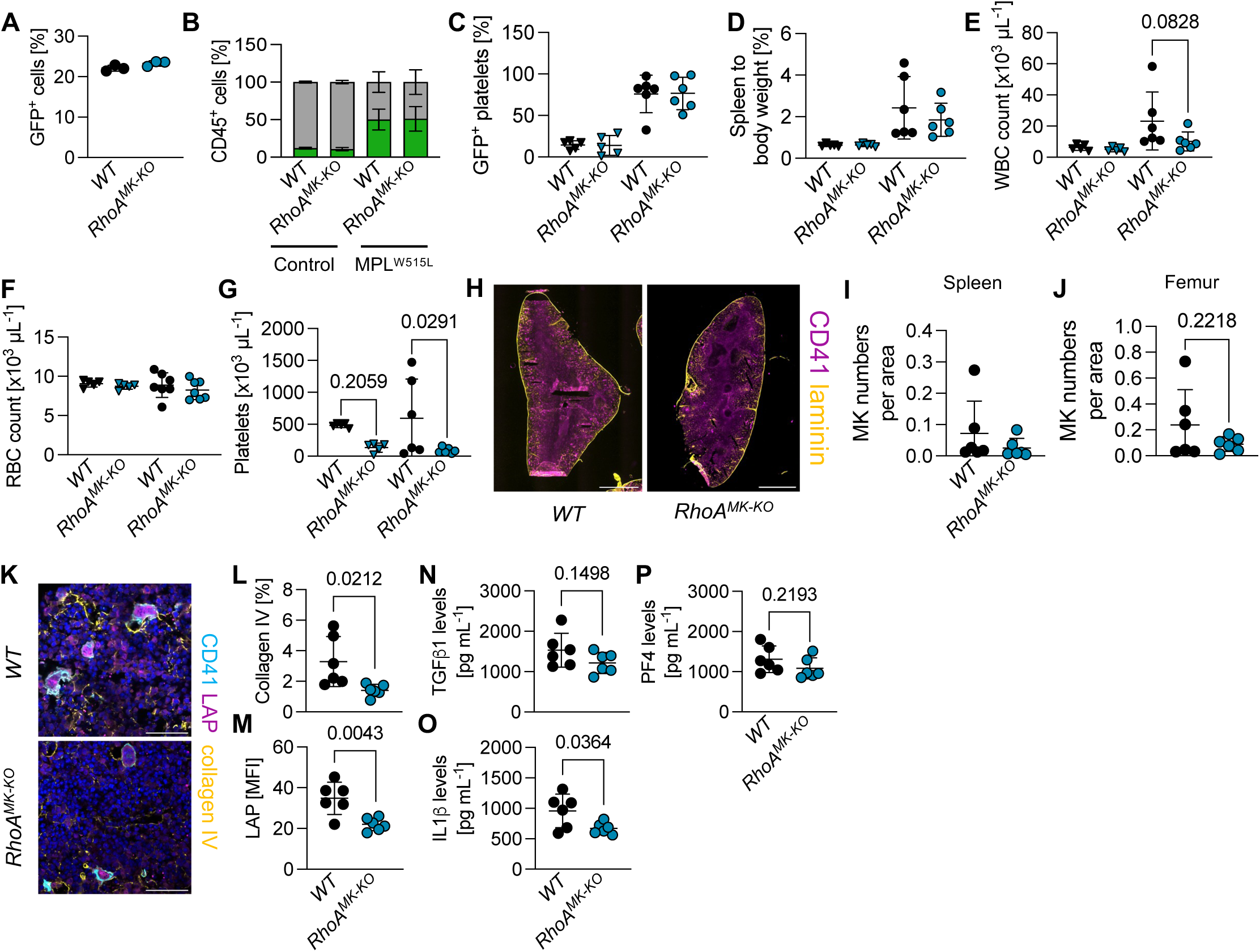
Attenuated fibrosis upon *RhoA*-deficiency in MPL^W515L^ transplanted mice. **(A)** Transduction efficiency of CD117^+^ cells derived from *WT* and *RhoA^MK-KO^* mice transduced with MPL^W515L^ one day after transduction. n=3 mice. Multiple unpaired Student’s t-tests. **(B, C)** Percentage of EGFP^+^ CD45^+^ cells **(B)** and platelets **(C)** of *WT* and *RhoA^MK-KO^* mice transplanted with Control or MPL^W515L^-transduced cells four weeks after transplantation. n=5-6 mice. Two-way ANOVA with Sidak’s correction for multiple comparisons and unpaired, two-tailed Student’s t-test. **(D)** Spleen to body weight of *WT* and *RhoA^MK-KO^* mice transplanted with Control or MPL^W515L^-transduced cells. n=5-6 mice. One-way ANOVA with Sidak’s correction for multiple comparisons. **(E, F)** White blood cell (WBC) **(E)** and red blood cell (RBC) counts **(F)** of transplanted *WT* and *RhoA^MK-KO^* mice. n=5-6 mice. One-way ANOVA with Sidak’s correction for multiple comparisons. **(G)** Platelet counts of transplanted *WT* and *RhoA^MK-KO^* mice. n=5-6 mice. One-way ANOVA with Sidak’s correction for multiple comparisons. **(H, I)** Visualization **(H)** and quantification **(I)** of splenic MKs of MPL^W515L^-transplanted *WT* and *RhoA^MK-KO^* mice. n=6 mice. Scale bars: 500 µm. Unpaired, two-tailed Student’s t-test. **(J)** Quantification of MK numbers in femoral cryosections of *WT* and *RhoA^MK-KO^* mice transplanted with MPL^W515L^-transduced cells. n=6 mice. Unpaired, two-tailed Student’s t-test. **(K-M)** Visualization **(K)** and quantification of collagen IV **(L)** and intracellular LAP/TGFβ1 **(M)** and in transplanted *WT* and *RhoA^MK-KO^* mice. n=6. Unpaired, two-tailed Student’s t-test. Scale bars: 50 µm. **(N-P)** Analysis of TGFβ1 **(N)**, IL1β **(O)** and PF4 levels **(P**) in bone marrow fluid of MPL^W515L^-transplanted *WT* and *RhoA^MK-KO^* mice. n=6 mice. Unpaired, two-tailed Student’s t-test. All data are presented as mean ± SD.

### Treatment with autophagy or Rho kinase inhibitors ameliorates MK clustering and fibrosis in vivo

To test the therapeutic potential of targeting secretory autophagy downstream of RhoA, we next transplanted mice with MPL^W515L^-transduced cells, allowed engraftment for one week and treated them with either the autophagy inhibitor HQ (50 mg/kg) or the pan-ROCK inhibitor Y (30 mg/kg) (**Supplemental Figure 4A**) for two weeks. Treatment with HQ resulted in unaltered EGFP-positivity in CD45-positive peripheral blood cells as well as platelets (**Supplemental Figure 4B, C**). We also did not observe differences in spleen weights (**Supplemental Figure 4D**), white or red blood cell and platelet counts between PBS-and HQ-treated mice (**Supplemental Figure 4E-G**), suggesting overall similar disease progression between both groups. To assess the effect of HQ treatment on TGFβ1 secretion and subsequent fibrosis, we visualized CD41, TGFβ1/LAP and collagen I in femoral cryosections (**Supplemental Figure 4H**). Interestingly, we found significantly lower levels of LAP within MKs of HQ-treated mice (**Supplemental Figure 4I**), which correlated to markedly lower collagen I deposition in the bone marrow (**Supplemental Figure 4J**) implying reduced fibrosis. While MK numbers were not significantly different (**Supplemental Figure 4K**), we observed less MK clustering, which is a hallmark of disease in MF patients (*17*). Ultimately, we tested cytokine levels in the bone marrow of PBS- and HQ-treated mice, which revealed a trending reduction in TGFβ1 after HQ treatment (**Supplemental Figure 4L**), while IL1β and PF4 levels were unchanged (**Supplemental Figure 4M, N**). Overall, HQ mildly improved disease development by attenuating collagen deposition.

Selective deletion of *RhoA* from MKs and platelets reduced cytokine release *in vivo* and inhibition of the RhoA effector ROCK promoted polyploidization and MK maturation *in vitro* (*see Supplemental Figure 1A*), a phenomenon that was previously leveraged to attenuate MF progression in studies using inhibitors of Aurora kinase A that are currently being tested in clinical trials (*13, 14*). To synergize these findings, we next treated MPL^W515L^-transplanted mice with the pan-ROCK inhibitor Y. Similarly to HQ, we did not observe differences in EGFP-positivity between PBS control- and Y-treated animals (**Figure 6A**), however, the number of EGFP-positive platelets was lower in Y-treated mice compared to PBS controls (**Figure 6B**). Intriguingly, spleen weights of Y-treated mice were significantly reduced compared to PBS-treated mice (**Figure 6C**), implying ameliorated disease upon ROCK inhibition. White and red blood cell counts were comparable (**Figure 6D, E**), but platelet counts, IPF and MPV were lower in Y-treated mice as compared to PBS controls (**Figure 6F-H**). When visualizing TGFβ1 and collagen deposition in the bone marrow, we observed significantly reduced intracellular LAP within MKs in Y-treated animals (**Figure 6I**), which correlated to ameliorated fibrosis as visualized by collagen IV staining (**Figure 6J, K**). While MK numbers were not significantly different between PBS- and Y-treated mice (**Figure 6L**), similar to HQ, MK clustering was less abundant in femoral cryosections. Most importantly, Y treatment resulted in a significant reduction in bone marrow levels of TGFβ1 and IL1β (**Figure 6M, N**), correlating to alleviated fibrosis. In line with our hypothesis, PF4 levels in the bone marrow were similar between both groups (**Figure 6O**), highlighting that Y treatment specifically affected cytokine release via secretory autophagy. Overall, we identified a targetable pathway that attenuated disease progression in the MPL^W515L^ transplant model by reducing the autophagy-dependent release of TGFβ1 and IL1β and subsequent fibrosis.

**Figure 6.**
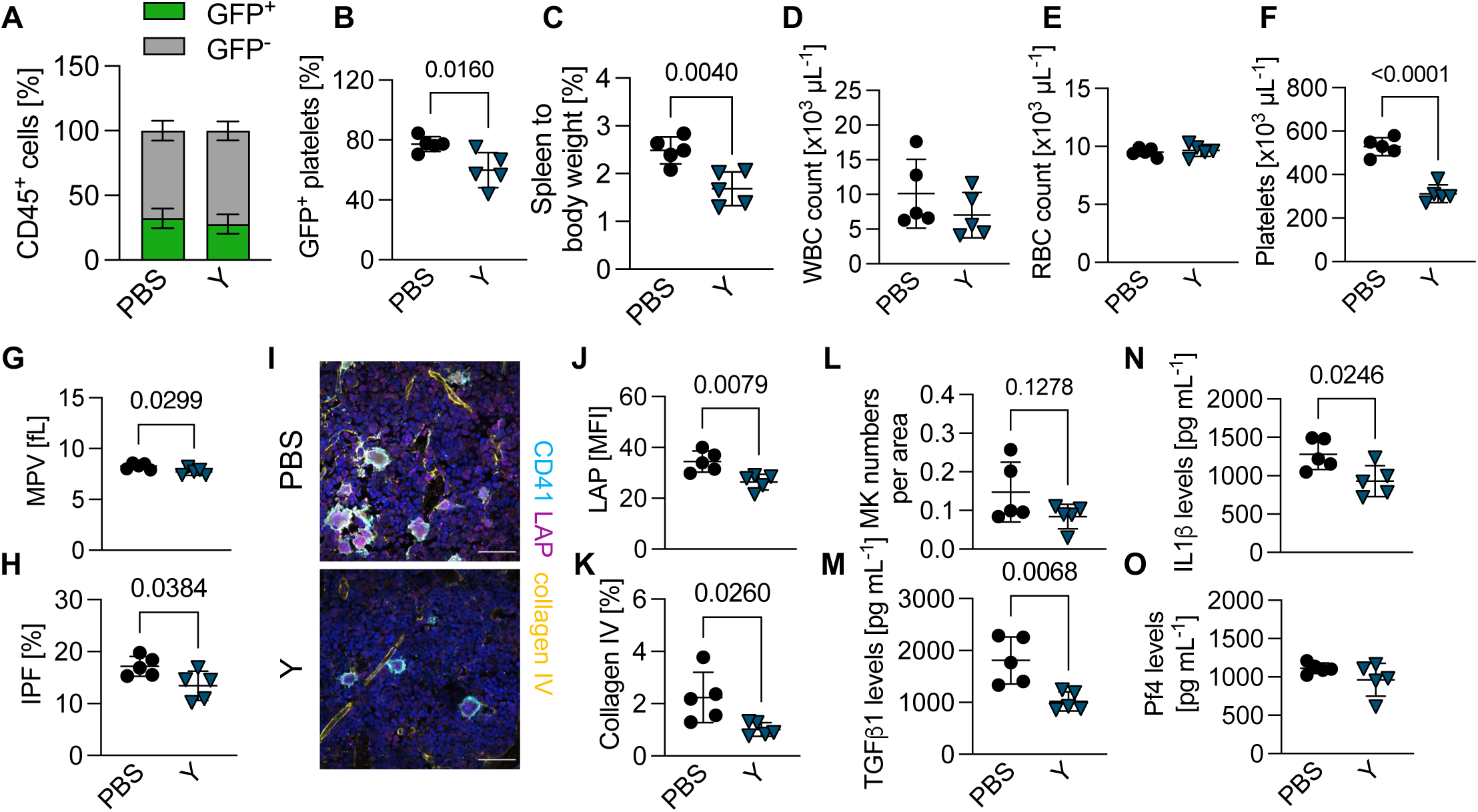
Autophagy/ROCK inhibition mitigates splenomegaly, cytokine burden and myelofibrosis. **(A, B)** Percentage of EGFP^+^ CD45^+^ cells **(A)** and platelets **(B)** of PBS- or Y27632 (Y)-treated mice transplanted with MPL^W515L^-transduced cells three weeks after transplantation. n=5 mice. Two-way ANOVA with Sidak’s correction for multiple comparisons and unpaired, two-tailed Student’s t-test. **(C)** Spleen to body weight of transplanted PBS- and Y-treated mice. n=5 mice. Unpaired, two-tailed Student’s t-test. **(D, E)** White blood cell (WBC) **(D)** and red blood cell (RBC) counts **(E)** of transplanted PBS- and Y-treated mice. n=5 mice. Unpaired, two-tailed Student’s t-test. **(F-H)** Platelet count **(F),** mean platelet volume (MPV) **(G)** and immature platelet fraction (IPF) **(H)** in transplanted PBS- and HQ-treated mice. n=5 mice. Unpaired, two-tailed Student’s t-test. All data are presented as mean ± SD. **(I-K)** Visualization **(I)** and quantification of intracellular LAP/TGFβ1 **(J)** and collagen IV **(K)** in transplanted PBS- and Y-treated mice. n=5. Unpaired, two-tailed Student’s t-test. Scale bars: 50 µm. **(L)** Quantification of MK numbers in femoral cryosections in transplanted PBS- and Y-treated mice. n=5 mice. Unpaired, two-tailed Student’s t-test. **(M-O)** Analysis of TGFβ1 **(M)**, IL1β **(N)** and PF4 levels **(O**) in bone marrow fluid of transplanted PBS- and Y-treated mice. n=5 mice. Unpaired, two-tailed Student’s t-test. All data are presented as mean ± SD. Schematic was generated using Biorender.com.

### Combination of autophagy/ROCK inhibitors with JAK2 inhibitors attenuates fibrosis progression beyond monotherapy

The current standard of care for the treatment of low-risk MF is the use of one of the approved JAK2 inhibitor (e.g. ruxolitinib), which significantly reduce spleen size and display partial disease-modifying qualities. However, the overall effects of JAK2 inhibition on cytokine burden and subsequent fibrosis are limited (*12, 58*). Our previous results suggested that autophagy/ROCK inhibition directly interferes with cytokine secretion and might thus represent a novel tool to suppress enhanced cytokine release in MF. To test this hypothesis *in vivo*, we treated MPL^W515L^ mice with the JAK2 inhibitor ruxolitinib (R) alone or in combination with HQ or Y for two weeks. Similar to single treatments with HQ and Y, no difference in EGFP positivity of CD45^+^ cells was observed (**Figure 7A**). Platelets of mice treated with R+HQ or R+Y had more variable EGFP-positivity, with some mice exhibiting a reduced amount of EGFP-positive platelets compared to both the vehicle and R monotherapy (**Figure 7B**). As expected, R treatment significantly reduced spleen weights in MPL^W515L^-transplanted mice, a phenomenon that was even more pronounced upon combination treatment with HQ or Y (**Figure 7C**). R similarly reduced leukocytosis and was even more potent in combination with HQ or Y (**Figure 7D**). While red blood cell counts were comparable between all groups, platelets tended to be lower in all treated groups, which was also reflected in a lower IPF (**Figure 7F, G**). When visualizing MKs and collagen deposition in the bone marrow by immunofluorescence, we did not observe differences in MKs counts between the groups (**Figure 7H, I**). However, collagen deposition was markedly attenuated only in mice treated with R+Y compared to R monotherapy, with a trending decrease upon R+HQ (**Figure 7J**). This was also reflected in a significant reduction in TGFβ1 levels in mice treated with R+Y and trending decreases in R+HQ compared to R monotherapy (**Figure 7K**). Similar to single treatment with HQ and Y, only the combination with the ROCK inhibitor Y mildly affected IL1β release (**Figure 7L**). Ultimately, we did not observe differences in PF4 levels in either of the groups, thus again emphasizing that HQ/Y treatment selectively affected secretory autophagy. In summary, we identified a so-far unknown secretion pathway of TGFβ1 and IL1β from MKs, which is distinct from conventional granule secretion. Moreover, we revealed that this pathway can be therapeutically leveraged to reduce cytokine burden and fibrosis in a preclinical mouse model of MF.

**Figure 7.**
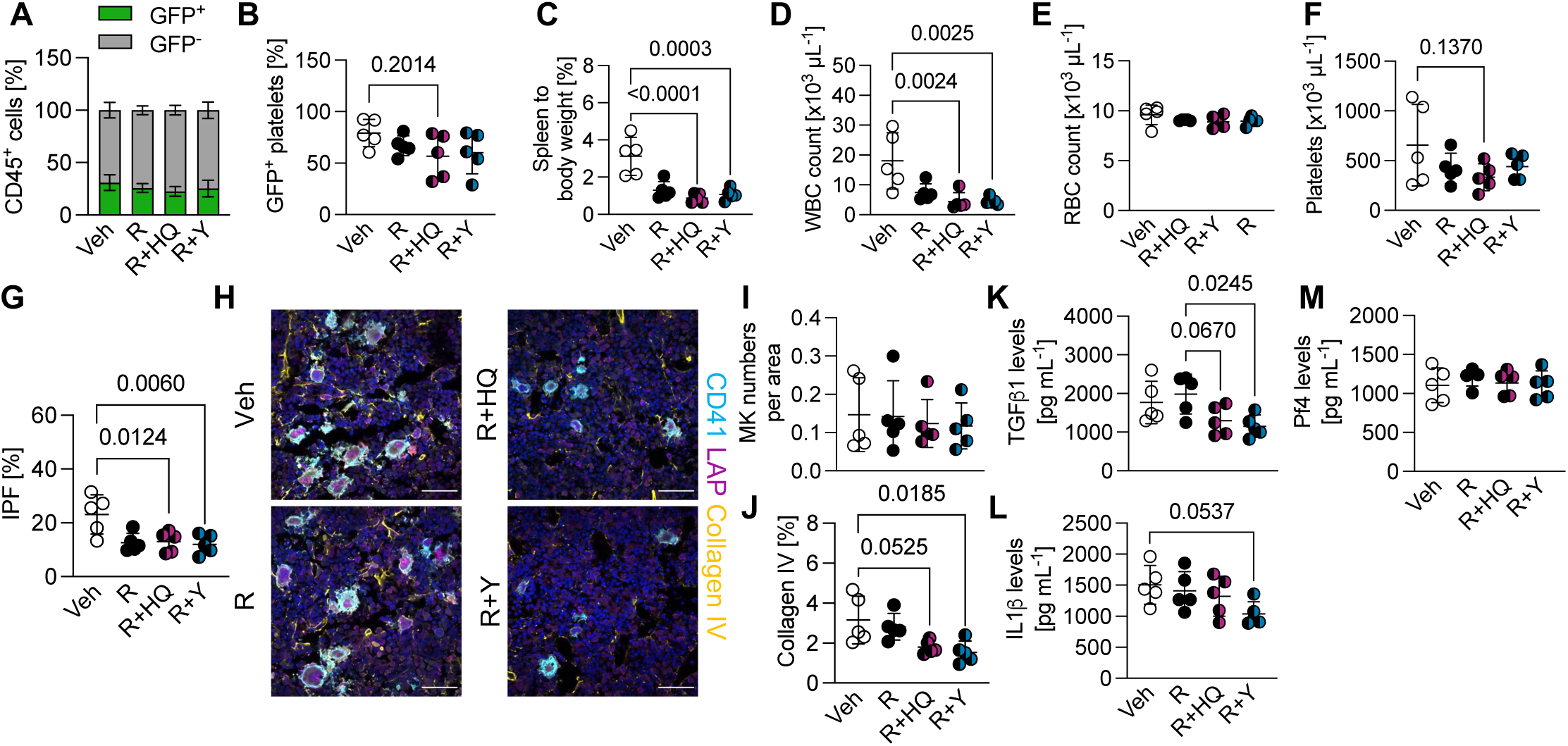
Combination therapy of autophagy/ROCK inhibitors with ruxolitinib alleviates fibrosis beyond monotherapy. **(A, B)** Percentage of EGFP^+^ CD45^+^ cells **(A)** and platelets **(B)** of vehicle (Veh; carboxymethylcellulose), ruxolitinib (R)-, ruxolitinib and hydroxychloroquine (HQ) (R+HQ)- and ruxolitinib and Y27632 (R+Y)-treated mice transplanted with MPL^W515L^-transduced cells three weeks after transplantation. n=5 mice. Two-way ANOVA with Sidak’s correction for multiple comparisons and one-way ANOVA with Sidak’s correction for multiple comparisons. **(C)** Spleen to body weight of transplanted Veh-, R-, R+HQ- and R+Y-treated mice. n=5 mice. One-way ANOVA with Sidak’s correction for multiple comparisons. **(D, E)** White blood cell (WBC) **(D)** and red blood cell (RBC) counts **(E)** of transplanted Veh-, R-, R+HQ- and R+Y-treated mice. n=5 mice. One-way ANOVA with Sidak’s correction for multiple comparisons. **(F, G)** Platelet count **(F)** and immature platelet fraction (IPF) **(G)** in transplanted Veh-, R-, R+HQ- and R+Y-treated mice. n=5 mice. One-way ANOVA with Sidak’s correction for multiple comparisons. **(H-J)** Visualization **(H)** and quantification of MK numbers **(I)** and collagen IV deposition **(J)** in transplanted Veh-, R-, R+HQ- and R+Y-treated mice. n=5 mice. One-way ANOVA with Sidak’s correction for multiple comparisons. Scale bars: 50 µm. **(K-M)** Analysis of TGFβ1 **(K)**, IL1β **(L)** and PF4 levels **(M**) in bone marrow fluid of transplanted Veh-, R-, R+HQ- and R+Y-treated mice. n=5 mice. One-way ANOVA with Sidak’s correction for multiple comparisons. All data are presented as mean ± SD.

## Discussion

Identifying novel pathways for the treatment of MF independent of JAK inhibition is urgently needed to improve patient care. Here, we demonstrate that MK-derived TGFβ1 is released via an unconventional secretion mechanism involving the autophagy machinery, which is controlled by the activity of RhoA and its downstream effector ROCK. Our data reveals that the pathologic release of TGFβ1 can be targeted by interfering with RhoA/ROCK-mediated secretory autophagy resulting in alleviated disease burden in a preclinical mouse model of MF. This represents a novel therapeutic target for MF.

While neoplastic transformation systemically affects a plethora of cell types that jointly drive disease progression, MKs play a crucial part in promoting fibrosis. Early studies revealed a tight association between atypical, dysmorphic MKs and increased extracellular matrix deposition in MF (*17, 59*). MKs derived from MF patients cultured *in vitro* exhibited reduced ploidy and increased cytokine release (*17*), in line with a reduced expression of the transcription factor GATA1, which is essential for MK differentiation (*60–62*). Due to their cell size and abundance of granules present within their cytoplasm, MKs are considered important regulators of paracrine and autocrine signaling in the bone marrow. At steady state, MKs regulate HSC maintenance through the secretion of CXCL4/PF4 and TGFβ1 (*4, 5*). Importantly, both cytokines have been implicated as major contributors to the progression of bone marrow fibrosis. MKs have been extensively demonstrated as the main source of TGFβ1 in MF (*17, 63, 64*). Using transmission electron microscopy and immunogold labeling, it was previously suggested that TGFβ1 was stored within MK α-granules (*35*), however, the specific increase in TGFβ1 secretion from dysmorphic MKs in MF suggests a distinct release mechanism.

Our data provides the first evidence for a role of secretory autophagy in regulating TGFβ1 secretion from MKs, inhibition of which reduces its release *in vitro* and *in vivo*. Furthermore, we identified increased basal autophagy in platelets of MPL^W515L^-transplaned mice, suggesting upregulation of autophagy upon neoplastic transformation. It was previously implied that autophagy inhibition enhances sensitivity of myeloproliferative clones towards ruxolitinib-induced apoptosis (*65*). Mutations in autophagy-related genes were also associated with worsened survival in MPN patients (*66, 67*), suggesting potential therapeutic options of autophagy inhibition even beyond cytokine secretion. This is reflected in our findings using *Atg5*-deficient cells in the MPL^W515L^ transplant model, in which disease progression was significantly attenuated resulting not only in lower TGFβ1 and IL1β secretion but also reduced clonal expansion (Figure 4). Our results using the autophagy inhibitor HQ reflect the reduction in cytokine secretion (Supplemental Figure 4), however, no differences in general disease onset were observed, suggesting that future studies should focus on identifying more specific inhibitors of autophagy.

Previous studies suggested secretory autophagy to mediate cytokine secretion from fibroblasts (*31*) with RhoA/ROCK1 signaling being identified as a main regulator of this process (*32, 52*). Interestingly, upregulation of the RhoA pathway in murine MKs (*33*), as well as RhoA GEFs in bone marrow MKs of MF patients were recently identified by scRNA Seq (*18, 34*), suggesting an important role of the GTPase in MKs in MF. Both deletion of *Rhoa* from the MK lineage as well as inhibition of ROCK signaling using the general ROCK inhibitor Y27632 significantly prevented fibrosis progression in an MPL^W515L^ transplant model of MF. Importantly, we observed an amelioration of general disease outcome even beyond decreased TGFβ1 secretion upon treatment with Y27632, as evident by a reduction in spleen size, platelet counts and IPF (Figure 6). A variety of MF therapeutic strategies aim to prevent MK dysplasia and defective polyploidization by targeting the cell cycle. One very promising protein target currently being tested in clinical trials is Aurora kinase A, inhibition of which increased MK maturation *in vitro* and thus overcame and prevented MK clustering and fibrosis in mouse models (*13, 14*). We observed a similar increase in polyploidization and MK maturation upon ROCK inhibition, suggesting that its targeting might concomitantly increase MK maturation and reduce cytokine secretion. In addition to TGFβ1, several other cytokines have been implicated in promoting MF progression. Recently, several groups reported an upregulation of the pro-inflammatory cytokine IL1β in MF patients and in a Jak2^V617F^ mouse model (*50, 56*). Notably, IL1β is another cytokine known to be released via secretory autophagy (*29, 30*), and we observed decreased IL1β levels upon RhoA/ROCK inhibition similar to TGFβ1, suggesting synergetic therapeutic benefits of targeting autophagy on both TGFβ1 and IL1β secretion. Of note, CXCL4 expression also positively correlates with disease severity in MF and deletion of the gene ameliorates fibrosis in preclinical mouse models (*68*). Mechanistically, CXCL4 appears to be internalized by stromal cells inducing a profibrotic transformation, inhibition of which similarly alleviates myelofibrosis in mice (*33*). While our data demonstrated unaltered PF4/CXCL4 levels upon autophagy/ROCK inhibition, we still observed protection from fibrosis, suggesting that targeting TGFβ1 and IL1β was sufficient in reducing collagen deposition in the MPL^W515L^ model.

The concept of targeting autophagy alongside conventional chemotherapies is not novel and a variety of clinical trials are evaluating beneficial effects of HQ in the treatment of patients with solid tumors (*69*). Our preclinical data suggests that targeting autophagy by inhibiting ROCKs would even further improve disease outcome in MPN patients. ROCK inhibitors are increasingly gaining attention as potential novel therapeutics for a variety of diseases, and the Food and Drug Administration recently approved the ROCK2 inhibitor Belumosudil for the treatment of Graft-versus-host-disease, implying manageable side effects of ROCK inhibition in patients (*70*). One drawback that the present study does not address is the relative contribution of the different ROCK isotypes to MF progression, which would be necessary to ultimately identify or develop inhibitors selectively targeting one ROCK isoform. Moreover, it would be interesting to assess whether ROCK inhibition can reverse already established fibrosis in mouse models. In summary, our results demonstrate a role of RhoA/ROCK-mediated secretory autophagy in mediating the release of TGFβ1 and IL1β from MKs independent of conventional granule secretion. We propose further investigation into either direct autophagy or ROCK inhibitors as novel therapeutic approaches for the treatment of MF, which may improve patient outcome beyond ruxolitinib monotherapy.

## Acknowledgements

We thank Ethan Walsey for excellent technical support and Dr. Jonas Jutzi for invaluable discussions and proofreading of the manuscript. We also thank Prof. Kimberley Tolias (Baylor College of Medicine, Houston, Texas) for kindly providing *Rhoa^fl/fl^* mice.

## Funding

Walter Benjamin Fellowship, German Research Foundation: BE 7766/2-1 (ICB) NHLBI, K99/R00 Pathway to Independence: 1K99HL175037 (MNB)

National Institutes of Health R35HL161175 (JEI)

## Author contributions

Conceptualization: ICB

Methodology and investigation: ICB, MNB, JL, VC, APS, BAC, RAJS, HGR

Supervision: JEI

Writing - original draft: ICB

Writing - review & editing: ICB, MNB, VC, RAJS, HGR, JEI

## Data and materials availability

All data needed to evaluate the conclusions in the paper are present in the paper and/or the Supplementary Materials.

## References

1. N. L. Asquith et al., The bone marrow is the primary site of thrombopoiesis. Blood 143, 272–278 (2024).

2. L. J. Noetzli, S. L. French, K. R. Machlus, New Insights Into the Differentiation of Megakaryocytes From Hematopoietic Progenitors. Arterioscler Thromb Vasc Biol 39, 1288–1300 (2019).

3. J. N. Thon et al., Cytoskeletal mechanics of proplatelet maturation and platelet release. J Cell Biol 191, 861–874 (2010).

4. I. Bruns et al., Megakaryocytes regulate hematopoietic stem cell quiescence through CXCL4 secretion. Nat Med 20, 1315–1320 (2014).

5. M. Zhao et al., Megakaryocytes maintain homeostatic quiescence and promote post-injury regeneration of hematopoietic stem cells. Nat Med 20, 1321–1326 (2014).

6. S. Sun et al., Single-Cell Analysis of Ploidy and Transcriptome Reveals Functional and Spatial Divergency in Murine Megakaryopoiesis. Blood, (2021).

7. H. Wang et al., Decoding Human Megakaryocyte Development. Cell Stem Cell 28, 535–549 e538 (2021).

8. J. Wang et al., CXCR4(high) megakaryocytes regulate host-defense immunity against bacterial pathogens. Elife 11, (2022).

9. J. Kota, N. Caceres, S. N. Constantinescu, Aberrant signal transduction pathways in myeloproliferative neoplasms. Leukemia 22, 1828–1840 (2008).

10. A. Tefferi, A. Pardanani, Myeloproliferative Neoplasms: A Contemporary Review. JAMA Oncol 1, 97–105 (2015).

11. W. Vainchenker, R. Kralovics, Genetic basis and molecular pathophysiology of classical myeloproliferative neoplasms. Blood 129, 667–679 (2017).

12. C. N. Harrison et al., Long-term findings from COMFORT-II, a phase 3 study of ruxolitinib vs best available therapy for myelofibrosis. Leukemia 30, 1701–1707 (2016).

13. N. Gangat et al., Aurora Kinase A Inhibition Provides Clinical Benefit, Normalizes Megakaryocytes, and Reduces Bone Marrow Fibrosis in Patients with Myelofibrosis: A Phase I Trial. Clin Cancer Res 25, 4898–4906 (2019).

14. Q. J. Wen et al., Targeting megakaryocytic-induced fibrosis in myeloproliferative neoplasms by AURKA inhibition. Nat Med 21, 1473–1480 (2015).

15. C. C. Ponce, F. C. M. de Lourdes, S. S. Ihara, M. R. Silva, The relationship of the active and latent forms of TGF-beta1 with marrow fibrosis in essential thrombocythemia and primary myelofibrosis. Med Oncol 29, 2337–2344 (2012).

16. M. Zingariello et al., A novel interaction between megakaryocytes and activated fibrocytes increases TGF-beta bioavailability in the Gata1(low) mouse model of myelofibrosis. Am J Blood Res 5, 34–61 (2015).

17. S. O. Ciurea et al., Pivotal contributions of megakaryocytes to the biology of idiopathic myelofibrosis. Blood 110, 986–993 (2007).

18. S. H. Jung et al., Different inflammatory, fibrotic, and immunologic signatures between pre-fibrotic and overt primary myelofibrosis. Haematologica, (2024).

19. J. A. Guerrero, et al., Gray platelet syndrome: proinflammatory megakaryocytes and alpha-granule loss cause myelofibrosis and confer metastasis resistance in mice. Blood 124, 3624–3635 (2014).

20. M. Bender et al., Dynamin 2-dependent endocytosis is required for normal megakaryocyte development in mice. Blood 125, 1014–1024 (2015).

21. I. C. Becker et al., G6b-B regulates an essential step in megakaryocyte maturation. Blood Adv, (2022).

22. I. Hofmann et al., Congenital macrothrombocytopenia with focal myelofibrosis due to mutations in human G6b-B is rescued in humanized mice. Blood 132, 1399–1412 (2018).

23. D. Peng, M. Fu, M. Wang, Y. Wei, X. Wei, Targeting TGF-beta signal transduction for fibrosis and cancer therapy. Mol Cancer 21, 104 (2022).

24. H. Chagraoui et al., Prominent role of TGF-beta 1 in thrombopoietin-induced myelofibrosis in mice. Blood 100, 3495–3503 (2002).

25. J. C. Yao et al., TGF-beta signaling in myeloproliferative neoplasms contributes to myelofibrosis without disrupting the hematopoietic niche. J Clin Invest 132, (2022).

26. J. Mascarenhas et al., A Phase Ib Trial of AVID200, a TGFbeta 1/3 Trap, in Patients with Myelofibrosis. Clin Cancer Res 29, 3622–3632 (2023).

27. L. Varricchio, et al., TGFbeta1 protein trap AVID200 beneficially affects hematopoiesis and bone marrow fibrosis in myelofibrosis. JCI Insight, (2021).

28. M. Ponpuak et al., Secretory autophagy. Curr Opin Cell Biol 35, 106–116 (2015).

29. N. Dupont et al., Autophagy-based unconventional secretory pathway for extracellular delivery of IL-1beta. EMBO J 30, 4701–4711 (2011).

30. L. Iula et al., Autophagy Mediates Interleukin-1beta Secretion in Human Neutrophils. Front Immunol 9, 269 (2018).

31. J. Nuchel et al., TGFB1 is secreted through an unconventional pathway dependent on the autophagic machinery and cytoskeletal regulators. Autophagy 14, 465–486 (2018).

32. A. U. Gurkar et al., Identification of ROCK1 kinase as a critical regulator of Beclin1-mediated autophagy during metabolic stress. Nat Commun 4, 2189 (2013).

33. D. Capitanio et al., Proteomic screening identifies PF4/Cxcl4 as a critical driver of myelofibrosis. Leukemia, (2024).

34. R. Li et al., A proinflammatory stem cell niche drives myelofibrosis through a targetable galectin-1 axis. Sci Transl Med 16, eadj7552 (2024).

35. R. A. Fava et al., Synthesis of transforming growth factor-beta 1 by megakaryocytes and its localization to megakaryocyte and platelet alpha-granules. Blood 76, 1946–1955 (1990).

36. T. Heib, C. Gross, M. L. Muller, D. Stegner, I. Pleines, Isolation of murine bone marrow by centrifugation or flushing for the analysis of hematopoietic cells - a comparative study. Platelets, 1–7 (2020).

37. M. Spindler, K. Mott, H. Schulze, M. Bender, Rapid isolation of mature murine primary megakaryocytes by size exclusion via filtration. Platelets 34, 2192289 (2023).

38. E. M. Pietras et al., Functionally Distinct Subsets of Lineage-Biased Multipotent Progenitors Control Blood Production in Normal and Regenerative Conditions. Cell Stem Cell 17, 35–46 (2015).

39. C. J. Pronk et al., Elucidation of the phenotypic, functional, and molecular topography of a myeloerythroid progenitor cell hierarchy. Cell Stem Cell 1, 428–442 (2007).

40. T. Kawamoto, M. Shimizu, A method for preparing 2- to 50-micron-thick fresh-frozen sections of large samples and undecalcified hard tissues. Histochem Cell Biol 113, 331–339 (2000).

41. Y. Pikman et al., MPLW515L is a novel somatic activating mutation in myelofibrosis with myeloid metaplasia. PLoS Med 3, e270 (2006).

42. F. J. de Sauvage et al., Physiological regulation of early and late stages of megakaryocytopoiesis by thrombopoietin. J Exp Med 183, 651–656 (1996).

43. A. L. Gurney, K. Carver-Moore, F. J. de Sauvage, M. W. Moore, Thrombocytopenia in c-mpl-deficient mice. Science 265, 1445–1447 (1994).

44. J. T. Lykins et al., Serglycin controls megakaryocyte retention of platelet factor 4 and influences megakaryocyte fate in bone marrow. Blood Adv, (2024).

45. T. J. Melia, A. H. Lystad, A. Simonsen, Autophagosome biogenesis: From membrane growth to closure. J Cell Biol 219, (2020).

46. E. Donohue et al., Inhibition of autophagosome formation by the benzoporphyrin derivative verteporfin. J Biol Chem 286, 7290–7300 (2011).

47. P. M. P. Ferreira, R. W. R. Sousa, J. R. O. Ferreira, G. C. G. Militao, D. P. Bezerra, Chloroquine and hydroxychloroquine in antitumor therapies based on autophagy-related mechanisms. Pharmacol Res 168, 105582 (2021).

48. Q. Wang et al., Rapamycin and bafilomycin A1 alter autophagy and megakaryopoiesis. Platelets 28, 82–89 (2017).

49. H. Schwertz, E. A. Middleton, Autophagy and its consequences for platelet biology. Thromb Res 231, 170–181 (2023).

50. M. F. Rahman et al., Interleukin-1 contributes to clonal expansion and progression of bone marrow fibrosis in JAK2V617F-induced myeloproliferative neoplasm. Nat Commun 13, 5347 (2022).

51. Y. Aman et al., Autophagy in healthy aging and disease. Nat Aging 1, 634–650 (2021).

52. M. O. Aguilera, W. Beron, M. I. Colombo, The actin cytoskeleton participates in the early events of autophagosome formation upon starvation induced autophagy. Autophagy 8, 1590–1603 (2012).

53. M. P. Avanzi et al., Rho kinase inhibition drives megakaryocyte polyploidization and proplatelet formation through MYC and NFE2 downregulation. Br J Haematol 164, 867–876 (2014).

54. C. Zhang et al., Endosidin2 targets conserved exocyst complex subunit EXO70 to inhibit exocytosis. Proc Natl Acad Sci U S A 113, E41–50 (2016).

55. B. A. Chua et al., Hematopoietic stem cells preferentially traffic misfolded proteins to aggresomes and depend on aggrephagy to maintain protein homeostasis. Cell Stem Cell 30, 460–472 e466 (2023).

56. S. Rai et al., Inhibition of interleukin-1beta reduces myelofibrosis and osteosclerosis in mice with JAK2-V617F driven myeloproliferative neoplasm. Nat Commun 13, 5346 (2022).

57. I. Pleines et al., Megakaryocyte-specific RhoA deficiency causes macrothrombocytopenia and defective platelet activation in hemostasis and thrombosis. Blood 119, 1054–1063 (2012).

58. N. Pemmaraju et al., Ten years after ruxolitinib approval for myelofibrosis: a review of clinical efficacy. Leuk Lymphoma 64, 1063–1081 (2023).

59. M. Schino et al., Bone marrow megakaryocytic activation predicts fibrotic evolution of Philadelphia-negative myeloproliferative neoplasms. Haematologica 106, 3162–3169 (2021).

60. L. Gilles et al., Downregulation of GATA1 drives impaired hematopoiesis in primary myelofibrosis. J Clin Invest 127, 1316–1320 (2017).

61. A. M. Vannucchi et al., A pathobiologic pathway linking thrombopoietin, GATA-1, and TGF-beta1 in the development of myelofibrosis. Blood 105, 3493–3501 (2005).

62. A. M. Vannucchi et al., Abnormalities of GATA-1 in megakaryocytes from patients with idiopathic myelofibrosis. Am J Pathol 167, 849–858 (2005).

63. S. Badalucco et al., Involvement of TGFbeta1 in autocrine regulation of proplatelet formation in healthy subjects and patients with primary myelofibrosis. Haematologica 98, 514–517 (2013).

64. M. C. Martyre, H. Magdelenat, M. C. Bryckaert, C. Laine-Bidron, F. Calvo, Increased intraplatelet levels of platelet-derived growth factor and transforming growth factor-beta in patients with myelofibrosis with myeloid metaplasia. Br J Haematol 77, 80–86 (1991).

65. C. Courdy et al., Targeting PP2A-dependent autophagy enhances sensitivity to ruxolitinib in JAK2(V617F) myeloproliferative neoplasms. Blood Cancer J 13, 106 (2023).

66. J. A. Machado-Neto et al., Autophagy inhibition potentiates ruxolitinib-induced apoptosis in JAK2(V617F) cells. Invest New Drugs 38, 733–745 (2020).

67. V. Visconte et al., Complete mutational spectrum of the autophagy interactome: a novel class of tumor suppressor genes in myeloid neoplasms. Leukemia 31, 505–510 (2017).

68. H. F. E. Gleitz et al., Increased CXCL4 expression in hematopoietic cells links inflammation and progression of bone marrow fibrosis in MPN. Blood 136, 2051–2064 (2020).

69. K. N. Townsend et al., Autophagy inhibition in cancer therapy: metabolic considerations for antitumor immunity. Immunol Rev 249, 176–194 (2012).

70. J. Barcelo, R. Samain, V. Sanz-Moreno, Preclinical to clinical utility of ROCK inhibitors in cancer. Trends Cancer 9, 250–263 (2023).

