## Supplemental Figures for "Inhibition of RhoA-mediated secretory autophagy in megakaryocytes mitigates myelofibrosis in mice"

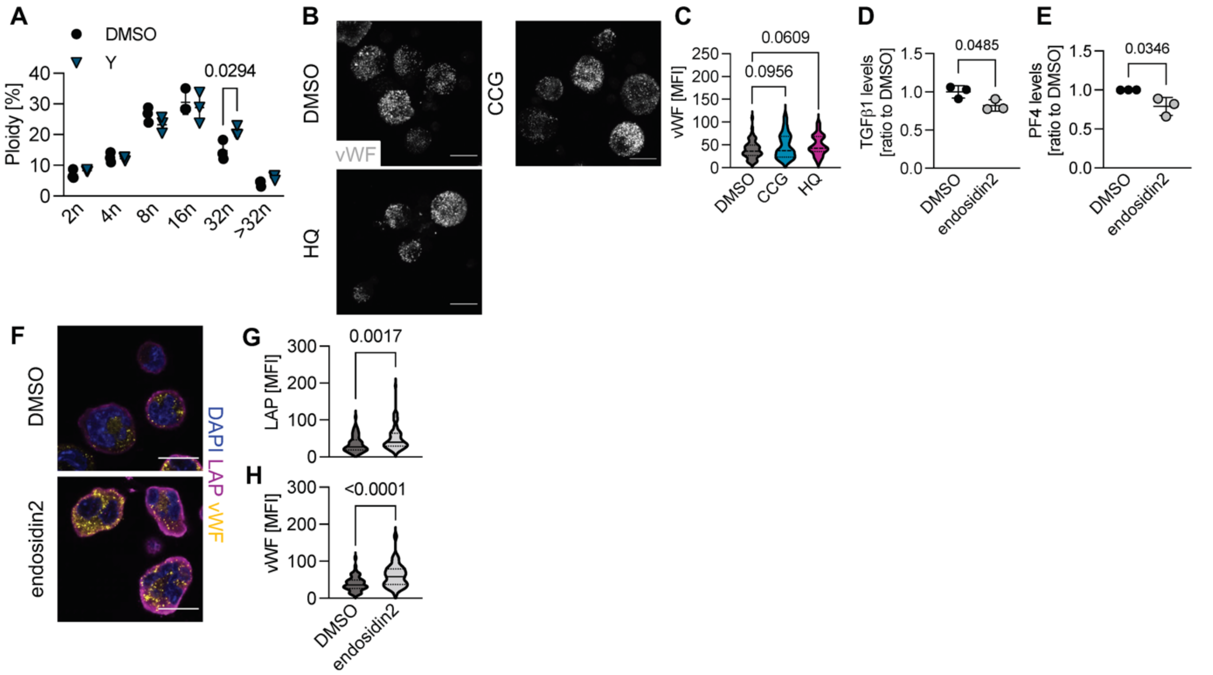


**Supplemental Figure 1. (A)** Ploidy analysis of *in vitro* matured megakaryocytes (MKs) differentiated in the presence of DMSO or 500 nM Y27632 (Y) for 72h. n=3 mice. Two-way ANOVA with Sidak's correction for multiple comparisons. **(B, C)** Visualization **(B)** and quantification **(C)** of vWF in enriched bone marrow-derived MKs treated with DMSO, 5 µM CCG or 5 µM HQ for 72h. Nuclei were counterstained using DAPI. Scale bars: 20 µm. At least 70 MKs were analyzed per condition. One-way ANOVA with Dunnett's test for multiple comparisons. **(D, E)** Analysis of transforming growth factor β1 (TGFβ1) **(D)** and platelet factor 4 (PF4) levels **(E)** in cell culture supernatant after treatment of hematopoietic stem and progenitor cells (HSPCs) with DMSO or 50 µM endosidin2 for 72h. n=3 mice. Unpaired, two-tailed Student's t-test. **(F-H)** Visualization **(F)** and quantification of mean fluorescence intensity (MFI) of TGFβ1/LAP **(G)** and von Willebrand Factor (vWF) **(H)** in enriched MKs after treatment with endosidin2 for 72h. n=3 mice. Unpaired, two-tailed Student's t-test. All data are presented as mean ± SD. All data are presented as mean ± SD. Schematics were generated using Biorender.com.

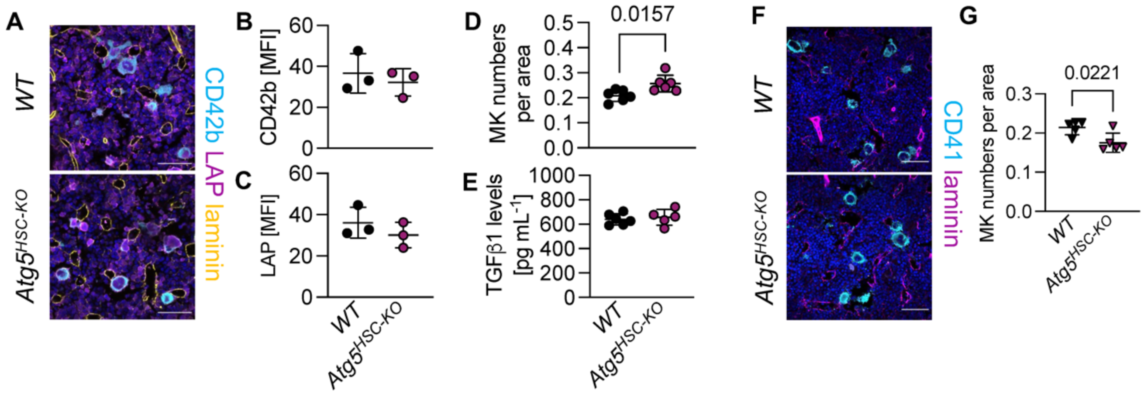


**Supplemental Figure 2. (A-C)** Visualization **(A)** and quantification of LAP/TGFβ1 **(B)**, the MK marker CD42b **(C)** and laminin in femoral cryosections of non-transplanted *WT* and *Atg5*$^{HSC\text{-}KO}$ mice. Nuclei were counterstained using DAPI. n=4 mice. Unpaired, two-tailed Student's t-test. Scale bars: 50 µm. **(D)** Quantification of MK numbers in femoral cryosections of *WT* and *Atg5*$^{HSC\text{-}KO}$ mice. n=6 mice. Unpaired, two-tailed Student's t-test. **(E)** Analysis of TGFβ1 levels in bone marrow fluid of *WT* and *Atg5*$^{HSC\text{-}KO}$ mice. n=6 mice. **(F, G)** Visualization **(F)** and quantification **(G)** of MK numbers in femoral cryosections of MSCV-EGFP mice transplanted with *WT* or *Atg5*$^{HSC\text{-}KO}$ cells four weeks after transplantation. n=5 mice. Unpaired, two-tailed Student's t-test.

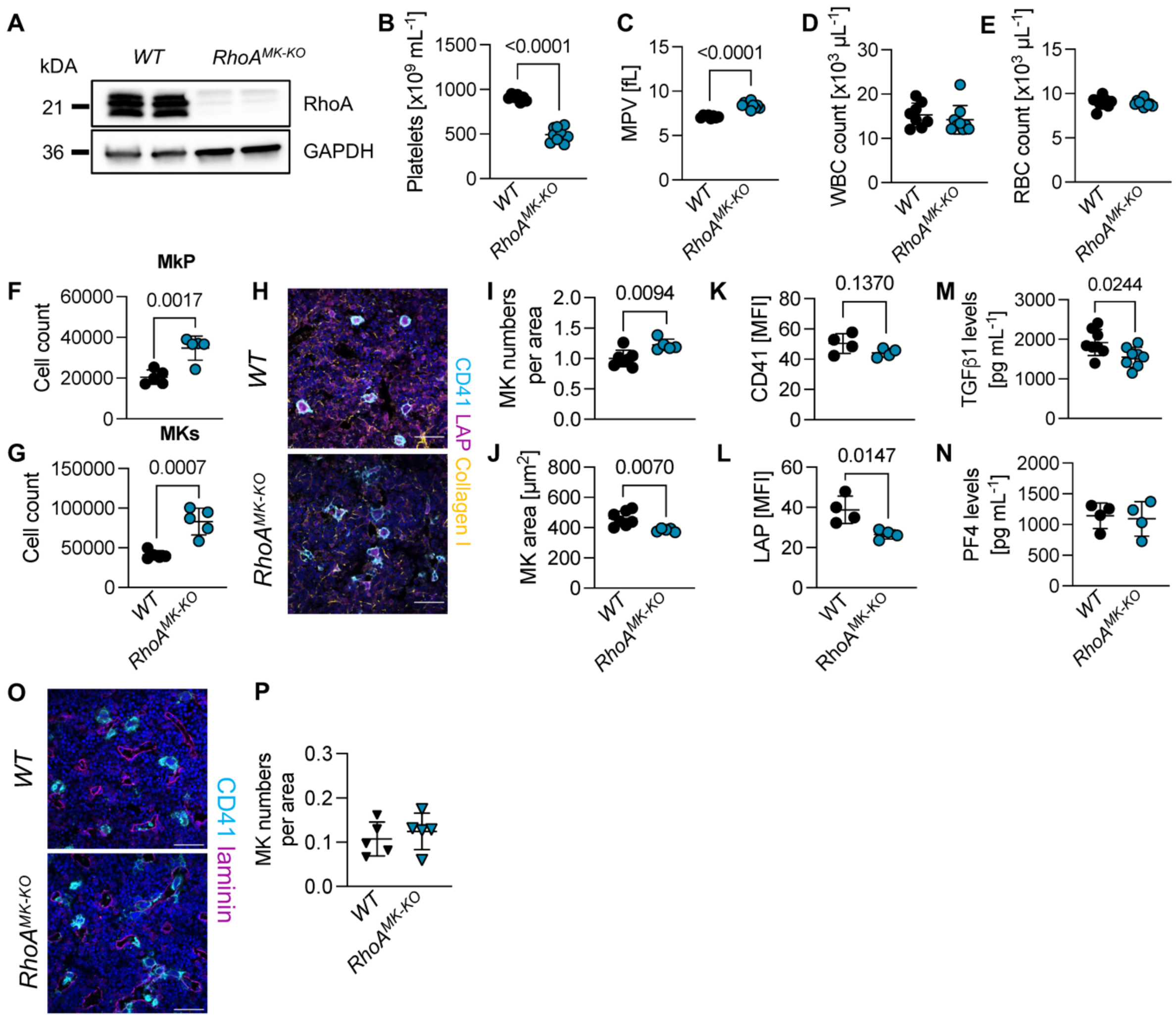


**Supplemental Figure 3. (A)** RhoA immunoblot of platelets derived from *WT* and *RhoA*$^{MK\text{-}KO}$. n=2 mice. **(B, C)** Platelet count **(B)** and mean platelet volume (MPV) **(C)** of *WT* and *RhoA*$^{MK\text{-}KO}$ mice. n=9 mice. Unpaired, two-tailed Student's t-test. **(D, E)** White blood cell (WBC) **(D)** and red blood cell (RBC) counts **(E)** of *WT* and *RhoA*$^{MK\text{-}KO}$ mice. n=9 mice. Unpaired, two-tailed Student's t-test. **(F, G)** Quantification of megakaryocyte progenitors (MkPs) **(F)** and MKs **(G)** in the bone marrow of *WT* and *RhoA*$^{MK\text{-}KO}$ mice by flow cytometry. n=5 mice. Unpaired, two-tailed Student's t-test. **(H-J)** Visualization **(H)** and quantification of MK numbers **(I)** and area **(J)** in femoral cryosections. Nuclei were counterstained using DAPI. n=6 mice. Unpaired, two-tailed Student's t-test. Scale bars: 50 µm. **(K, L)** Quantification of CD41 **(K)** and LAP/TGFβ1 in MKs **(L)** in femoral cryosections of non-transplanted *WT* and *RhoA*$^{MK\text{-}KO}$ mice. n=4 mice. Unpaired, two-tailed Student's t-test. **(D)** Quantification of MK numbers in femoral cryosections of *WT* and *RhoA*$^{MK\text{-}KO}$ mice. n=6 mice. Unpaired, two-tailed Student's t-test. **(M, N)** Analysis of TGFβ1 **(M)** and PF4 levels **(N)** in bone marrow fluid of *WT* and *RhoA*$^{MK\text{-}KO}$ mice. n=4-8 mice. **(O, P)** Visualization **(O)** and quantification **(P)** of MK numbers in femoral cryosections of MSCV-EGFP mice transplanted with *WT* or *RhoA*$^{MK\text{-}KO}$ cells four weeks after transplantation. n=5 mice. Unpaired, two-tailed Student's t-test.

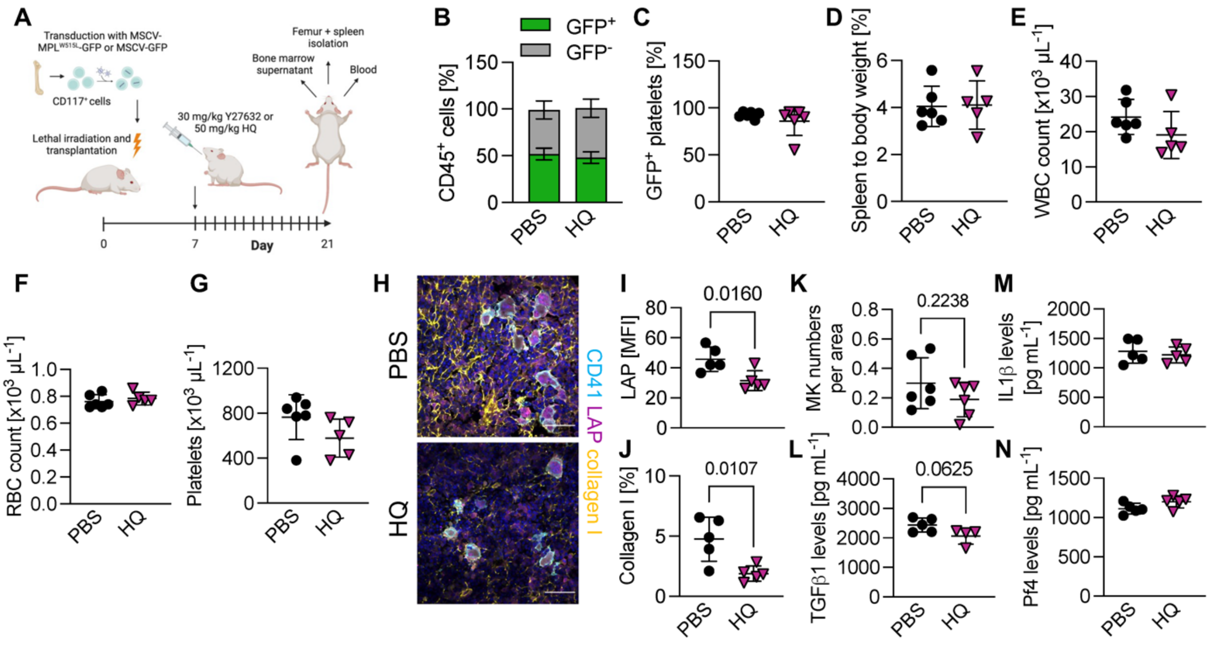


**Supplemental Figure 4. (A)** Schematic of MPL$^{W515L}$ transplant model and treatment regimen in BALB/cJ mice. **(B, C)** Percentage of EGFP$^{+}$ CD45$^{+}$ cells **(B)** and platelets **(C)** of PBS or hydroxychloroquine (HQ)-treated mice transplanted with MPL$^{W515L}$-transduced cells three weeks after transplantation. n=5 mice. Two-way ANOVA with Sidak's correction for multiple comparisons and unpaired, two-tailed Student's t-test. **(D)** Spleen to body weight of transplanted PBS- and HQ-treated mice. n=5. Unpaired, two-tailed Student's t-test. **(E, F)** White blood cell (WBC) **(E)** and red blood cell (RBC) counts **(F)** of transplanted PBS- and HQ-treated mice. n=5 mice. Unpaired, two-tailed Student's t-test. **(G)** Platelet count in transplanted PBS- and HQ-treated mice. n=5 mice. Unpaired, two-tailed Student's t-test. **(H-J)** Visualization **(H)** and quantification of intracellular LAP/TGFβ1 **(I)** and collagen I deposition **(J)** in transplanted PBS- and HQ-treated mice. At least 30 MKs and four FOVs were analyzed per mouse. n=5 mice. Unpaired, two-tailed Student's t-test. Scale bars: 50 µm. **(K)** Quantification of MK numbers in femoral cryosections in transplanted PBS- and HQ-treated mice. n=5 mice. Unpaired, two-tailed Student's t-test. **(L-N)** Analysis of TGFβ1 **(L)**, interleukin 1β (IL1β) **(M)** and PF4 levels **(N)** in bone marrow fluid of transplanted PBS- and HQ-treated mice. n=5 mice. Unpaired, two-tailed Student's t-test.
